# Variability in porcine microRNA genes and its association with mRNA expression and lipid phenotypes

**DOI:** 10.1101/2020.04.17.038315

**Authors:** Emilio Mármol-Sanchez, María Gracia Luigi-Sierra, Anna Castelló, Dailu Guan, Raquel Quintanilla, Raul Tonda, Marcel Amills

## Abstract

**Background:** Mature microRNAs (miRNAs) play an important role in repressing the expression of a wide range of mRNAs. The variability of miRNA genes and their corresponding 3’UTR binding sites might disrupt canonical conserved miRNA-mRNA pairing, thus modifying gene expression patterns. The presence of polymorphic sites in miRNA genes and their association with gene expression phenotypes and complex traits has been poorly characterized in pigs so far.

**Results:** By analyzing whole-genome sequences from 120 pigs and wild boars from Europe and Asia, we have identified 285 single nucleotide polymorphisms (SNPs) mapping to miRNA loci, as well as 109,724 SNPs located in predicted 7mer-m8 miRNA binding sites within porcine 3’UTRs. Porcine miRNA genes show a reduced SNP density compared with their flanking non-miRNA regions. By sequencing the genomes of 5 Duroc boars, we have identified 12 miRNA SNPs that have been subsequently genotyped in their offspring (N = 345, Lipgen population). Association analyses between miRNA SNPs and 38 lipid-related traits as well as hepatic and muscle microarray expression phenotypes recorded in the Lipgen population were carried out. The most relevant association detected was the one between the genotype of the rs319154814 (G/A) SNP located in the apical loop of the ssc-miR-326 hairpin precursor and *PPP1CC* mRNA levels in the liver (*q*-value = 0.058). This result was subsequently confirmed by qPCR (*P*-value = 0.027). The rs319154814 (G/A) genotype was also associated with several fatty acid composition traits.

**Conclusions:** Porcine miRNA genes show a reduced variability consistent with strong purifying selection, particularly in the seed region, which plays a critical role in miRNA binding. Although it is generally assumed that SNPs mapping to the seed region are the ones with the strongest consequences on mRNA expression, we show that a SNP mapping to the apical region of ssc-miR-326 is significantly associated with the hepatic mRNA levels of the *PPP1CC* gene, one of its predicted targets. Although experimental confirmation of such interaction has been obtained in humans but not in pigs, this result highlights the need of further investigating the functional effects of miRNA polymorphisms located outside the seed region on gene expression in pigs.

## Background

Mature microRNA transcripts (miRNAs) are short (∼22 nt) non-coding RNAs which play an essential role in the regulation of gene expression [1]. During the biogenesis of miRNAs, one strand of the miRNA duplex binds to the guide-strand channel of an Argonaute protein forming a miRNA-induced silencing complex (miRISC) with the ability to repress mRNA expression through binding to specific 3’UTR target sites [2]. In postembryonic cells, this repressor mechanism mainly acts by destabilizing the mRNA through decapping and poly(A)-tail shortening [3, 4] and less often by hindering translation [2]. The binding of the miRNA to its 3’UTR target site depends critically on the sequence of the seed region, which encompasses nucleotides (nt) 2-8 from the 5’ end of the mature miRNA and interacts with the target site through Watson-Crick pairing [1]. Polymorphisms located within the seed region might affect the proper recognition of mRNA targets [5, 6]. Nevertheless, imperfect seed matches can be compensated by nucleotides 13^th^-to-16^th^ of the mature miRNA, thus providing additional anchor pairing to the seed region [2]. Other sites relevant for miRNA processing and function are the basal UG, flanking CNNC and apical UGU motifs [7], as well as a mismatched GHG motif [8], all of which contribute to facilitate miRNA processing [2]. Variability in the 3’UTR miRNA binding sites can also affect gene expression at the post-transcriptional level [5,9,10].

The evolutionary conservation of miRNAs across species is associated with their levels of expression [11, 12] as well as with the functional importance of the regulatory networks they modulate [13, 14]. Besides, the conservation of the repertoire of mRNAs targeted by a given miRNA depends mostly on the age of the associated miRNA gene, with novel miRNAs acquiring targets more rapidly than ancient miRNAs [15]. Conserved sites are particularly enriched in the 5’end of 3’UTR [12], although selective pressure on miRNA binding sites is not uniform across miRNA genes [16].

Saunders et al. (2007) [5] investigated the variability of 474 human miRNA genes and found that the single nucleotide polymorphism (SNP) density is lower in miRNA loci when compared to their flanking regions. Moreover, Saunders et al. (2007) [5] found that ∼90% of human miRNA genes do not contain polymorphisms, and the majority of SNPs mapping to miRNAs are located outside the seed region, thus evidencing that the variability of this critical functional motif evolves under strong selective constraints [5]. Indeed, polymorphisms in the first 14 nucleotides of the mature miRNA, and particularly those within the seed region, might abrogate the binding of the miRNA to its 3’UTR targets, leading to an extensive rewiring of the miRNA-mediated regulatory network [17]. Mammalian miRNA knockouts often display abnormal phenotypes, reduced viability and clinical disorders [2], although functional redundancy amongst miRNA family members might mitigate to some extent the severity of such manifestations [2]. Polymorphisms within miRNA loci laying outside the seed region can also affect the processing and stability of miRNAs during their maturation and loading into the functional silencing complex [18, 19]. Moreover, Saunders et al. (2007) [5] showed that a broad array of predicted miRNA target sites in the 3’UTR of mRNAs are polymorphic, a finding that suggests that purifying selection on these regions is less intense than in miRNA genes [17]. Overall, these findings suggest functional conserved mechanisms of miRNA-mediated gene regulation influenced by polymorphic sites at both miRNA loci and their 3’UTR binding sites [20].

The wild ancestors of pigs were independently domesticated in the Near East and China 10,000 YBP [21]. Domestic pigs then spread worldwide, becoming one of the most important sources of animal protein for humans and diversifying in an extensive array of breeds with distinct morphological and productive features [22]. Pig phenotypes might be explained, at least partially, by modifications in the microRNA-mediated regulation of gene expression [23]. Indeed, several studies have reported associations between SNPs in miRNA binding sites and porcine phenotypic variation [24–29], while fewer studies have investigated the association between SNPs in miRNA genes and complex traits in pigs [30–33]. For instance, Chai et al. (2018) reported a significant causal relationship between a SNP in the seed of ssc-miR-378-3p and extensive modifications of its target mRNA repertoire [33]. Moreover, the authors associated the presence of the observed miRNA seed variant to a structural reconfiguration of the hairpin, leading to an enhanced expression of the mature miRNA [33].

The polymorphism of miRNA genes has not been systematically characterized in pigs despite its potential impact on gene regulation and phenotypic variation. In the current work, we aimed to elucidate the patterns of variability of miRNA genes in European and Asian wild boars and domestic pigs as well as to infer whether such patterns are influenced by purifying selection. Moreover, we wanted to infer whether SNPs in miRNA genes are associated with liver and muscle gene expression traits and lipid phenotypes recorded in a Duroc pig population.

## Methods

### Characterizing the polymorphisms of miRNA genes and their 3’UTR binding sites in a worldwide sample of pigs and wild boars

#### Retrieval of Porcine Whole-Genome Sequences

Whole-genome sequences (WGS) from 120 wild boars and domestic pigs (*Sus scrofa*) were retrieved from the NCBI Sequence Read Archive (SRA) database (https://www.ncbi.nlm.nih.gov/sra). Most of these WGS have been reported in previous publications [34–37] and detailed information can be found in **Additional file 1: Table S1**. The 120 WGS corresponded to Asian domestic pigs (ADM, N = 40), Asian wild boars (AWB, N = 20), European domestic pigs (EDM, N = 40) and European wild boars (EWB, N = 20). The ADM group included WGS from Meishan, Tongchen, Jinhua, Rongchan, Wuzhishan, Tibetan, Sichuan, Hetao, Minzhu, Bamaixang and Laiwu pigs, while the EDM group was composed of WGS from Pietrain, Mangalitza, Iberian, Duroc, American Yucatan (from America, but with a European origin), Yorkshire, Landrace, Hampshire and Large-white pigs. Both AWB and EWB samples were selected according to their geographic distribution (**Additional file 1: Table S1**).

The EWB group contained one sample from the Near East. Raw data in SRA format were downloaded from SRA public repositories and converted into fastq format by using the fastq-dump 2.8.2 tool available in the SRA-toolkit package (ncbi.github.io/sra-tools/).

#### Whole-genome sequence data processing and calling of single nucleotide polymorphisms

Fastq paired-end files generated from SRA data were filtered according to their quality, and sequence adapters were trimmed by making use of the Trimmomatic software (v.0.36) with default parameters [38]. Trimmed paired-end sequences were aligned against the *Sus scrofa* reference genome (Sscrofa11.1) [39] with the BWA-MEM algorithm [40] and default settings. Sequence alignment map (SAM) formatted files were sorted and transformed into binary (BAM) formatted files. PCR duplicates were subsequently identified with the *MarkDuplicates* package from the Picard tool (https://broadinstitute.github.io/picard/) and removed to perform INDEL realignment with the *IndelRealigner* package from the Genome Analysis Toolkit (GATK v.3.8) [41]. Base quality score recalibration (BQSR) was implemented with the *BaseRecalibrator* package from GATK v.3.8. Variant calling was implemented with the *HaplotypeCaller* function according to GATK best practices [41]. Individual gVCF formatted files, including both polymorphic and homozygous blocks, were generated with the *GenotypeGVCFs* package from the GATK v.3.8 tool [41], and they were merged into separate multi-individual variant calling format (VCF) files containing single polymorphic and INDEL sites for each defined population (i.e. ADM, EDM, AWB and EWB). Finally, variants were distributed into SNP and INDEL files with the

*SelectVariants* package and quality filtered using the *FilterVariants* function from GATK v.3.8 [41]. For SNPs, we used the following parameters: QD < 2.0, FS > 60.0, MQ < 40.0, MQRankSum < -12.5, ReadPosRankSum < -8.0, SOR > 3.0. INDELs were not taken into consideration in the present study because INDEL detection based on commonly used pipelines for variant calling have shown low reliability and reproducibility across tools and analyzed datasets [42, 43].

#### Identification of single nucleotide polymorphisms in miRNA genes and in their 3’UTR binding sites

Single nucleotide polymorphisms mapping to annotated porcine miRNA loci (N = 370) were retrieved by making use of the curated Sscrofa11.1 annotation for miRNA regions available in the miRCarta v1.1 [44] and miRBase [45] databases. Additionally, annotated mature miRNA loci (N = 409) within miRNA genes (N = 370) were retrieved, and SNPs located in the mature and seed regions (2^nd^ to 8^th^ positions at 5’ end in the mature miRNA) were identified. A comprehensive description of miRNA gene regions and miRNA-mRNA interactions can be found in **Fig. 1A** and **1B**, respectively.

**Fig. 1:**
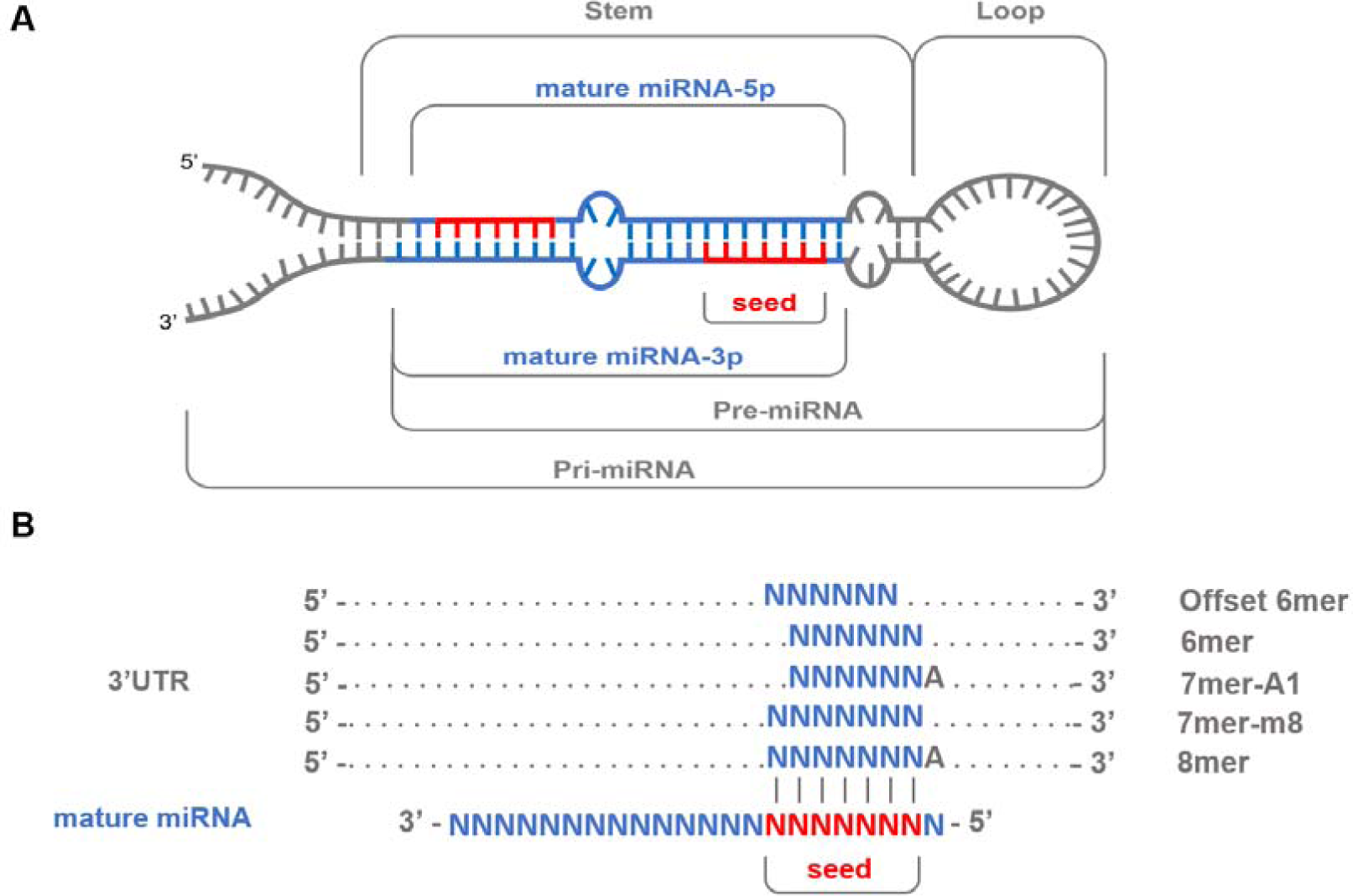
Schematic representation of (**A**) precursor miRNA sequence hairpin conformation, with mature miRNAs located in the stems (higlighted in blue) and their corresponding seed regions (in red). (**B**) Canonical miRNA-mRNA interactions where each binding site is conformed by 6-7 Watson-Crick pairings (vertical lines) to the seed region of the mature miRNA (2^nd^ to 8^th^ 5’ nucleotides). As reported elsewhere [2,47,48], 7mer-m8 and 8mer binding sites are considered to be the most functionally active canonical miRNA-mRNA interactions.

We also retrieved the annotated 3’UTR porcine mRNA transcripts from Ensembl repositories (v.100, http://www.ensembl.org/info/data/ftp/index.html) and the corresponding set of sequences was interrogated against the seed regions of the 409 porcine mature miRNAs considered in our study. Seed sequences were reverse-complemented and searched along the 3’UTR sequences of mRNA genes by making use of the *locate* function of the SeqKit toolkit [46]. All 7mer-m8 canonical seed pairing sites, as well as 8mer seed pairing sites (**Fig. 1**), were identified for each miRNA seed, and, based on such information, the corresponding putative miRNA-mRNA target pairs were established. Additional non-canonical miRNA-mRNA interactions were not considered (**Fig. 1B**) as they are regarded as less biologically significant than 7mer-m8 or 8mer interactions [2,47,48]. Subsequently, the genomic location of both 7mer-m8 and 8mer matching regions were determined, and SNPs residing in such predicted miRNA binding sites were gathered.

Principal components analyses (PCA) were performed with the smartPCA software [49]. These PCA analyses were based on either (1) autosomal whole-genome SNPs, (2) autosomal SNPs from miRNA genes, (3) autosomal SNPs from 3’UTRs, or (4) autosomal SNPs from 8mer and 7mer-m8 3’UTR sites. When implementing the PCA based on whole-genome data, SNPs complying with the following parameters were retained: 1) minimum allele frequency > 0.05, and 2) Hardy-Weinberg equilibrium exact test *P*-value > 0.001. In contrast, no filtering was performed in the PCA based on miRNA SNPs because there are a few hundreds.

#### Frequency and distribution of SNPs in miRNA genes

To investigate the distribution of SNPs along miRNA genes, SNPs were classified according to their location, i.e., SNPs within the seed region of mature miRNAs were flagged as “*seed*”, whereas SNPs located in mature miRNAs but outside the seed were classified as “*mature*”. Additionally, SNPs classified as “*mature*” were assigned to the following subtypes: 1) “*anchor*” (1^st^ position at 5’ end), and 2) “*supplemental pairing*” (13^th^ to 18^th^ position from 5’ end). The remaining SNPs were assigned to the “*precursor*” class. Allele frequencies were estimated in the whole set of porcine samples (N = 120), as well as in each EDM, ADM, EWB and AWB group independently (**Additional file 2: Table S2**).

In order to calculate the SNP density in precursor, mature and seed miRNA regions, we first calculated the total nucleotide length of each of these regions in the set of porcine miRNA loci. S*eed length* was calculated by considering 7 nucleotide positions in each of the annotated mature miRNAs (N = 409) mapping to the 370 miRNA loci considered in our study (7 bp × 409 = 2,863 bp). The total length of the 409 mature miRNAs was determined (∼22 bp × 409 = 8,916 bp), and the total *seed length* (2,863 bp) was subtracted from 8,916 bp to obtain the *mature length* (6,053 bp), which corresponds to the total length of all mature miRNA sequences excluding the seeds. The *precursor length* (22,229 bp) was calculated by subtracting the *seed length* (2,863 bp) and *mature length* (6,053 bp) from the total length (31,145 bp) of all miRNA loci (N = 370). Then, the SNP density (*D*) in each of these regions was calculated as follows:

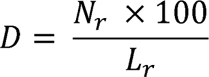

where *N_r_* is the total number of miRNA SNPs (N = 285), or the number of SNPs detected in each of the defined regions, (i.e., 221, 52 and 12 SNPs in precursor, mature and seed regions, respectively); *L_r_* is the total nucleotide length of all miRNA loci (31,145 nts), or the total number of nucleotides in miRNA loci belonging to the “*precursor*” (22,229 bp), “*mature*” (6,053 bp) or “*seed*” (2,863 bp) regions. These calculations would yield the number of SNPs per bp for each category. We decided to adjust this estimate to a window of 100 bp, which results in the multiplication of *N_r_* by 100 in the above formula. We also compared the SNP density between miRNA loci (N = 370) and their flanking genomic regions (± 1 kb) by considering 10 upstream and downstream nucleotide bins comprising 100 bp each. Differences in SNP density across miRNA loci, as well as compared with surrounding regions, were estimated with a non-parametric approach using a Mann Whitney U test [50].

### Investigating the association of miRNA polymorphisms with gene expression and phenotype data recorded in Duroc pigs

#### Whole-Genome sequencing of five Duroc pigs

In the year 2,003, five Duroc boars were selected and used as founders of a half-sib population of purebred Duroc pigs devoted to the production of high-quality cured ham. By whole-genome sequencing of these 5 individuals, we aimed to identify miRNA SNPs and investigate their association with gene expression traits and lipid phenotypes recorded in the offspring of the 5 boars (Lipgen population, N = 345). Genomic DNA was extracted [51] and sequenced at the Centre Nacional d’Anàlisi Genòmica (CNAG, Barcelona, Spain). Paired-end multiplex libraries were prepared according to the instructions of the manufacturer with the KAPA PE Library Preparation kit (Kapa Biosystems, Wilmington, MA). Libraries were loaded to Illumina flow-cells for cluster generation prior to producing 100 bp paired-end reads on a HiSeq2000 instrument following the Illumina protocol. Base calling and quality control analyses were performed with the Illumina RTA sequence analysis pipeline according to the instructions of the manufacturer. Quality-checked filtered reads were mapped to the *Sus scrofa* genome version 11.1 and processed for SNP calling according to GATK best practices recommendations [41] and the same protocol defined in a previous section (please see the section describing Whole-genome sequence data processing and calling of single nucleotide polymorphisms). Since only five boars were sequenced, SNPs within miRNA loci were retrieved without quality filtering criteria, in an effort to maximize the number of identified miRNA SNPs.

#### Recording of phenotypes related with gene expression and lipid traits in the Lipgen population

A total of 350 Duroc barrows, sired by the five Duroc founder boars mentioned before were used as a resource population (Lipgen population [52, 53]) to evaluate associations between miRNA SNPs and phenotypes related with gene expression and lipid traits. The five sequenced boars were mated with 400 sows in three different farms and one offspring per litter was selected for phenotypic recording (only 350 individuals provided valid records). All selected piglets were weaned, castrated and subsequently fattened in four contemporary batches at the IRTA pig experimental farm in Monells (Girona, Spain) under intensive standard conditions. Once they reached ∼122 kg of live weight (∼190 days of age), they were slaughtered in a commercial abattoir following the guidelines of current Spanish legislation (https://www.boe.es/buscar/doc.php?id=BOE-A-1995-3942). After slaughtering, tissue samples from *gluteus medius* (GM) and *longissimus dorsi* (LD) skeletal muscles and liver were obtained according to previously described methods [54–56].

Total DNA was extracted from each sample following Vidal et al. 2005 [51]. A subset of 345 DNA samples from the initial set of 350 pigs were successfully obtained and processed for further genotyping. Total RNA was extracted from GM (N = 89 pigs) and liver (N = 87 pigs) tissue samples by using the Ribopure isolation kit (Ambion, Austin, TX). Expression mRNA profiles were characterized by hybridizing total RNA to GeneChip porcine arrays (Affymetrix Inc., Santa Clara, CA), which encompass a total of 23,998 probes [54, 57]. Further details about tissue collection, sample selection, RNA isolation and microarray hybridization procedures can be found in Cánovas et al. (2010) [54]. Microarray data pre-processing, background correction, normalization and log_2_-transformation of expression estimates were performed with a robust multi-array average method per probe [58]. The *mas5calls* function from the affy R package [58] was then applied to identify probes displaying intensity signals above the background noise. This function applies a Wilcoxon signed rank-based gene expression presence/absence detection algorithm for labeling expressed probes in each sample. Control probes and those with expression levels below the detection threshold in more than 50% of samples were discarded from further analyses. By doing so, we only retained probes corresponding to mRNAs that were robustly expressed in GM muscle (12,131 probes) and liver (12,787 probes) tissues. Probes from GeneChip Porcine Genomic array identifiers (Affymetrix, Inc., Santa Clara, CA) of GM skeletal muscle and liver tissues were assigned to mRNA genes annotated in the Sscrofa11.1 assembly [39] using the BioMart tool [59].

With regard to lipid-related phenotypes, 38 traits were measured in the Duroc Lipgen population, i.e. backfat thickness measured between 3^rd^ and 4^th^ ribs, backfat thickness at the last rib, ham fat thickness, and intramuscular fat content and composition of GM and LD skeletal muscle samples (N = 345) [53, 55] (**Additional file 3: Table S3**). Briefly, intramuscular fatty acid (IMF) content in the GM and LD muscles was measured with the Near Infrared Transmittance technique (NIT, Infratec 1625, Tecator Hoganas, Sweden), while muscle cholesterol measurements were inferred following Cayuela et al. [60]. Gas chromatography of methyl esters was used to determine muscle fatty acids (FA) composition, including the percentages of saturated (SFA), unsaturated (UFA), monounsaturated (MUFA) and polyunsaturated (PUFA) fatty acids. Live and carcass weights (used as covariates in the statistical analyses), as well as backfat and ham fat thickness, were measured on a regular basis prior (live weight and backfat thickness) and after slaughter (carcass weight and ham fat thickness). Mean and standard deviations of lipid traits recorded the Lipgen population (N = 345) are described in **Additional file 3: Table S3**.

#### Genotyping of a panel of single nucleotide polymorphisms mapping to microRNA genes in the Lipgen population

Twelve SNPs mapping to miRNA loci and segregating in the five sequenced parental Duroc boars were genotyped in the 345 Duroc pigs from the Lipgen population (**Table 1**). Briefly, selected miRNA SNPs and their flanking regions (60 upstream and downstream bp) were used for assay design with the Custom TaqMan Assay Design Tool website (https://www5.appliedbiosystems.com/tools/cadt/; Life Technologies). Genotyping tasks were carried out at the Servei Veterinari de Genètica Molecular of the Universitat Autònoma of Barcelona (http://sct.uab.cat/svgm/en) by using a QuantStudio 12K Flex Real-Time PCR System (Thermo Fisher Scientific, Barcelona, Spain).

**Table 1:**
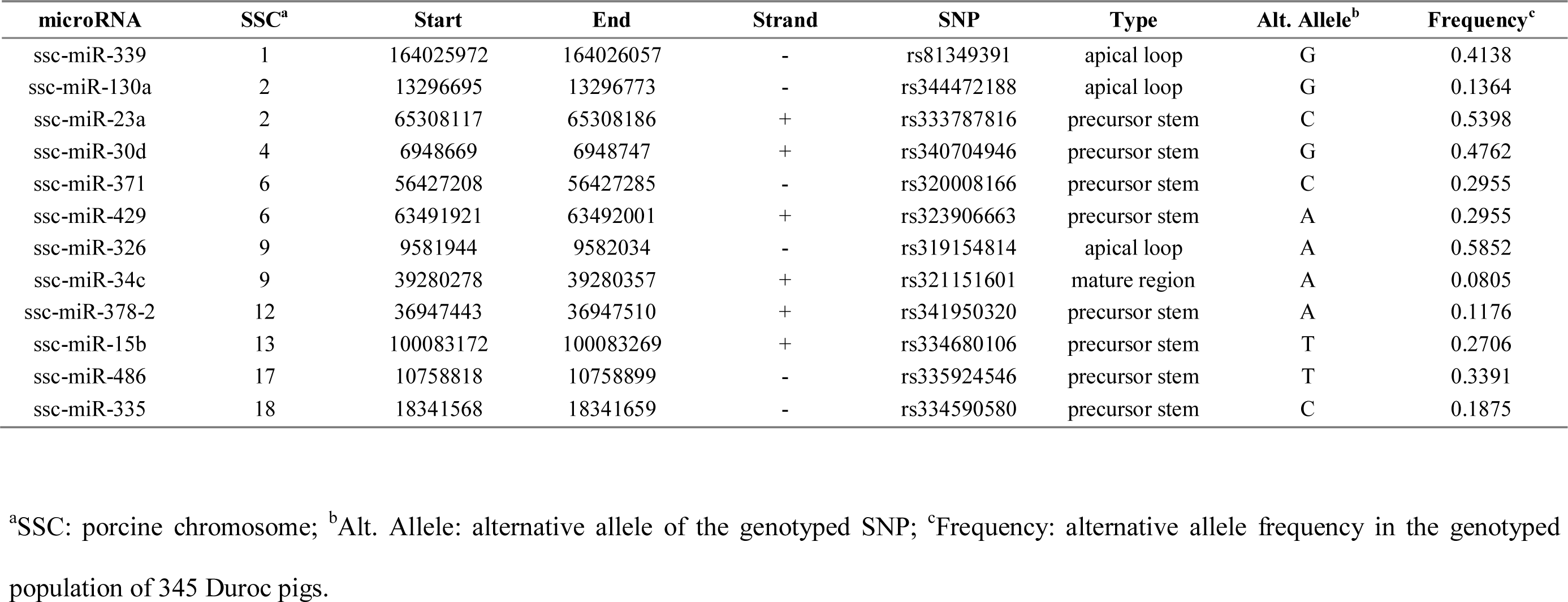
List of miRNA polymorphisms genotyped in a population of Duroc pigs (N = 345).

| microRNA | SSC<sup>a</sup> | Start | End | Strand | SNP | Type | Alt. Allele<sup>b</sup> | Frequency<sup>c</sup> |
| --- | --- | --- | --- | --- | --- | --- | --- | --- |
| ssc-miR-339 | 1 | 164025972 | 164026057 | - | rs81349391 | apical loop | G | 0.4138 |
| ssc-miR-130a | 2 | 13296695 | 13296773 | - | rs344472188 | apical loop | G | 0.1364 |
| ssc-miR-23a | 2 | 65308117 | 65308186 | + | rs333787816 | precursor stem | C | 0.5398 |
| ssc-miR-30d | 4 | 6948669 | 6948747 | + | rs340704946 | precursor stem | G | 0.4762 |
| ssc-miR-371 | 6 | 56427208 | 56427285 | - | rs320008166 | precursor stem | C | 0.2955 |
| ssc-miR-429 | 6 | 63491921 | 63492001 | + | rs323906663 | precursor stem | A | 0.2955 |
| ssc-miR-326 | 9 | 9581944 | 9582034 | - | rs319154814 | apical loop | A | 0.5852 |
| ssc-miR-34c | 9 | 39280278 | 39280357 | + | rs321151601 | mature region | A | 0.0805 |
| ssc-miR-378-2 | 12 | 36947443 | 36947510 | + | rs341950320 | precursor stem | A | 0.1176 |
| ssc-miR-15b | 13 | 100083172 | 100083269 | + | rs334680106 | precursor stem | T | 0.2706 |
| ssc-miR-486 | 17 | 10758818 | 10758899 | - | rs335924546 | precursor stem | T | 0.3391 |
| ssc-miR-335 | 18 | 18341568 | 18341659 | - | rs334590580 | precursor stem | C | 0.1875 |

<sup>a</sup>SSC: porcine chromosome; <sup>b</sup>Alt. Allele: alternative allele of the genotyped SNP; <sup>c</sup>Frequency: alternative allele frequency in the genotyped population of 345 Duroc pigs.

#### Association analyses between miRNA SNPs and mRNA expression and lipid phenotypes

Genotype data corresponding to the 12 miRNA SNPs mentioned before (**Table 1**) were processed with the PLINK software [61] in order to generate formatted files for subsequent analyses. The Genome-Wide Efficient Mixed-Model Association (GEMMA) software [62] was used to implement association analyses between genotyped SNPs and phenotypes related with microarray gene expression data and lipid traits. The following univariate mixed model was used:

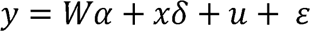

Where y is the phenotypic vector of recorded phenotypes for each individual; α is a vector indicating the intercept plus the considered fixed effects, i.e., batch effect with 4 categories (all traits), farm of origin effect with 3 categories (all traits) and laboratory of processing with 2 categories (GM and liver microarray expression data). The α vector also includes the following covariates: IMF of LD (for LD fatty acid α composition traits), IMF of GM (for GM fatty acid composition traits), live weight (for backfat thickness) and carcass weight (for ham fat thickness); *W* corresponds to the incidence matrix relating phenotypes with their corresponding effects; *x* is the genotype vector for the 12 miRNA polymorphisms; δ is the allele substitution effect for each δ polymorphism; *u* is a vector of random individual genetic effects with an n-dimensional multivariate normal distribution MVN_n_ (0, λ τ K), where τ corresponds to the variance of the residual errors, λ is the ratio between the two variance components and K is the known relatedness matrix derived from SNP information; and residual errors. is the vector of ε Association analyses were performed between each miRNA polymorphism (**Table 1**) and the 38 lipid-related traits listed in **Additional file 3: Table S3**. Moreover, we explored the associations between miRNA SNPs and expression levels of mRNAs that fulfilled the following conditions: 1) the 3’UTR of the mRNA contains at least one 7mer-m8 site complementary to the seed of the miRNA harboring the SNP to be tested (as predicted with the *locate* tool from SeqKit software [46]), 2) the existence of experimentally validated mRNA-miRNA interactions between the given mRNA and the tested miRNA have been confirmed in humans according to information provided in the DIANA-Tarbase v8 database [63]. The first condition was established because we were interested in focusing our analysis on miRNA-mRNA pairs for which a direct physical interaction (through complementarity between the seed of the miRNA and a target site in the 3’UTR of the mRNA) is feasible. The second condition was imposed because there is extensive evidence that in silico prediction or miRNA binding sites can have high false positive rates [64]. We considered as valid miRNA-mRNA interactions those experimentally validated in humans by using wet lab methods such as cross-linking, ligation and sequencing of hybrids (CLASH), photoactivatable ribonucleoside-enhanced and high-throughput sequencing of RNA isolated by crosslinking and immunoprecipitation (PAR-CLIP and HITS-CLIP) and luciferase assays [63].

Importantly, the seeds of the porcine miRNAs harboring the 12 genotyped SNPs were completely conserved with regard to the seeds of the human orthologous miRNA genes (**Additional file 4: Fig. S1**). This feature further supports the extrapolation of miRNA-mRNA interactions experimentally validated in humans [63] to pigs. For the sake of completeness, we also carried out a complementary association analysis between miRNA SNPs and the mRNA levels of the whole sets of genes expressed in the GM and liver tissues (i.e., without applying the two conditions mentioned before) but the results of such analyses should be interpreted with caution due to reasons stated above.

In association analyses with lipid traits or gene expression phenotypes, the alternative hypothesis H_1_ : δ ≠ 0 was contrasted against the null hypothesis H_0_ : δ = 0 with a likelihood ratio test. The statistical significance of the associations between miRNA SNPs and lipid and mRNA expression phenotypes was assessed with a false discovery rate (FDR) approach [65].

#### Pathway enrichment analysis

The lists of probes/genes significantly associated at the nominal level (*P*-value < 0.05) with miRNA SNPs genotypes (N = 12) after considering the criteria for miRNA target inference (see previous Methods section) were used as inputs for pathway enrichment analyses. The ClueGO v2.5.0 plug-in application [66] within the Cytoscape 3.8.2 software [67] was used for determining enriched Reactome and KEGG pathways. A one-sided hypergeometric test of significance was applied for determining enriched terms and multiple testing correction was implemented with a false discovery rate approach [65].

#### Confirming associations between ssc-miR-326 rs319154814 genotypes and gene expression data by RT-qPCR

The hepatic mRNA levels of three of the mRNA transcripts amongst the most significantly associated with ssc-miR-326 rs319154814 (G/A) genotype were analyzed by reverse transcription-quantitative polymerase chain reaction (RT-qPCR). The β-actin (*ACTB*) and TATA-Box binding protein (*TBP*) genes were used as endogenous controls, as previously reported [68]. In brief, 100 mg of liver tissue from 10 selected samples (5 from each GG and AA genotypes) were submerged in 1 mL TRIzol reagent (Invitrogen Corp., Carlsbad, CA, United States) and subsequently homogenized using the Lysing Matrix D reagent (MP Biomedicals, Santa Ana, CA) in a Precellys 24 tissue homogenizer (Bertin Instruments, Rockville, MD). Total RNA isolation was performed following the protocol described by Rio et al. (2010) [69]. Reverse transcription was achieved with the High-Capacity cDNA Reverse Transcription Kit (Applied Biosystems, Foster City, CA) following the instructions of the manufacturer. We used 1 μg of total RNA as a template in a final volume of 20 μl and the synthesized cDNA was diluted to 1:20 in Milli-Q water. One pair of primers spanning exon-exon junctions were designed for each gene (**Additional file 5: Table S4**) with the Primer Express software (Life Technologies Corporation). Real-time qPCR reactions contained 10 l of μ SYBR Select Master Mix, 300 nM of each primer, and 5 μl of 1:20 cDNA dilution, in a final volume of 20 μl. Reactions were run in an ABI PRISM 7900HT instrument (Applied Biosystems, Foster City, CA). The thermal cycle was set as follows: 2 min at 50 °C, one denaturing step at 95 °C during 10 min, followed by 40 cycles of 15 sec at 95 DC and 1 min at 60 °C. Moreover, a melting curve analysis was performed (i.e., 95 °C for 15 sec, 60 °C for 15 sec and a gradual increase in temperature, with a ramp rate of 1% up to 95 °C, followed by a final step of 95 °C for 15 sec), in order to assess the specificity of the reactions. A standard curve with serial dilutions from a pool including all the cDNA samples was used to assess whether amplification efficiencies of the three genes under analysis were comprised in the 90 to 110% range. All reactions were run in triplicate.

The 2^-Ct^ method [70] was used for the relative quantification of gene expression (Rq), using the group of GG samples as a calibrator. Subsequently, Rq values were log_2_ transformed and the significance of expression differences between GG and AA genotypes was assessed using Welch’s t-test for unpaired groups of samples [71] implemented in the *t.test* R function.

*Measuring the expression of the ssc-miR-326 gene in pigs with different rs319154814 genotypes*.

Liver samples from 9 AA and 9 GG pigs (rs319154814 genotype) were used to measure the expression of the ssc-miR-326 gene. Total RNA was isolated with TRIzol as previously described and reverse transcribed with the TaqMan Advanced MicroRNA Reverse Transcription Kit (Applied Biosystems, Foster City, CA), following the instructions of the manufacturer. Total RNA (10 ng) diluted in a volume of 2 μl, was used as a template to generate and pre-amplify cDNA by carrying out four consecutive reactions (poly-A tailing, adaptor ligation, reverse transcription and pre-amplification). The resulting pre-amplified cDNA was diluted to 1:50 in Milli-Q water. To measure ssc-miR-326 expression levels, both let-7a and miR-26a-5p were chosen as endogenous controls following Timoneda et al. (2012) [72]. To conduct the experiments for the three miRNAs under investigation (one target and two controls), available inventoried TaqMan advanced miRNA assays (Applied Biosystems, Foster City, CA) for each miRNA were purchased. The reactions were performed in a final volume of 15 μl that contained: 3.75 μl of 1:50 cDNA, 3 μl of nuclease-free water, 7.5 μl of TaqMan Fast Advanced Master Mix (2×), and 0.75 μl TaqMan Advanced miRNA Assay (20×). Each reaction was carried out in triplicate in a 384-well plate. Reactions were run in a QuantStudio 12K Flex Real-Time PCR System (Applied Biosystems, Foster City, CA). The thermal cycle program was the same used for qPCR experiments on mRNA transcripts. No standard dissociation curve was performed. The results were analyzed applying the 2^-Ct^ method [70] for the relative quantification of miRNA expression (Rq), using the group of GG samples as a calibrator. Subsequently, Rq values were log_2_ transformed and the significance of expression differences between GG and AA genotypes was assessed using a Welch’s t-test for unpaired groups [71].

## Results

### Differential segregation of SNPs mapping to microRNA genes in pigs and wild boars from Europe and Asia

A total of 58,537,491 million SNPs were identified with the GATK haplotype caller tool [41] in a data set comprising 120 WGS from 40 EDM, 40 ADM, 20 EWB and 20 AWB individuals (**Additional file 1: Table S1**). The distribution of these SNPs within different annotated regions (i.e. protein coding genes, exons, introns, 3’UTRs, miRNAs, lncRNAs, snRNAs, snoRNAs and pseudogenes) can be found in **Additional file 6: Table S5**. The majority of these SNPs were biallelic (96.82%), 777,008 (1.33%) were tri-allelic, 1,864,388 (3.18%) had a deletion allele and 57,878 (0.098%) showed four or more alleles (multi-allelic). From the set of multi-allelic SNPs, 1,765 SNPs mapped to exons, and 531 of them were clustered in a set of 74 genes enriched in olfactory receptors (**Additional file 7: Table S6**). This latter result is consistent with the potential presence of copy number variation (CNV) in such locations [73]. Alternative allele frequencies were consistently high (> 0.5) for 9.25% of variants, whereas low (between 0.05 and 0.01) and very low (< 0.01) alternative allele frequencies were detected in 27.85% and 19.62% of SNPs, respectively.

After filtering, 19,720,314 autosomal whole-genome SNPs were selected for assessing population structure based on PCA clustering techniques. The PCA showed a strong genetic differentiation amongst Asian and European populations (**Fig. 2A**). In contrast, domestic pigs and wild boars, and particularly those with European origin, did not show such stark divergence. Asian pigs and wild boars displayed some level of genetic differentiation and were more diverse than their European counterparts.

**Fig. 2:**
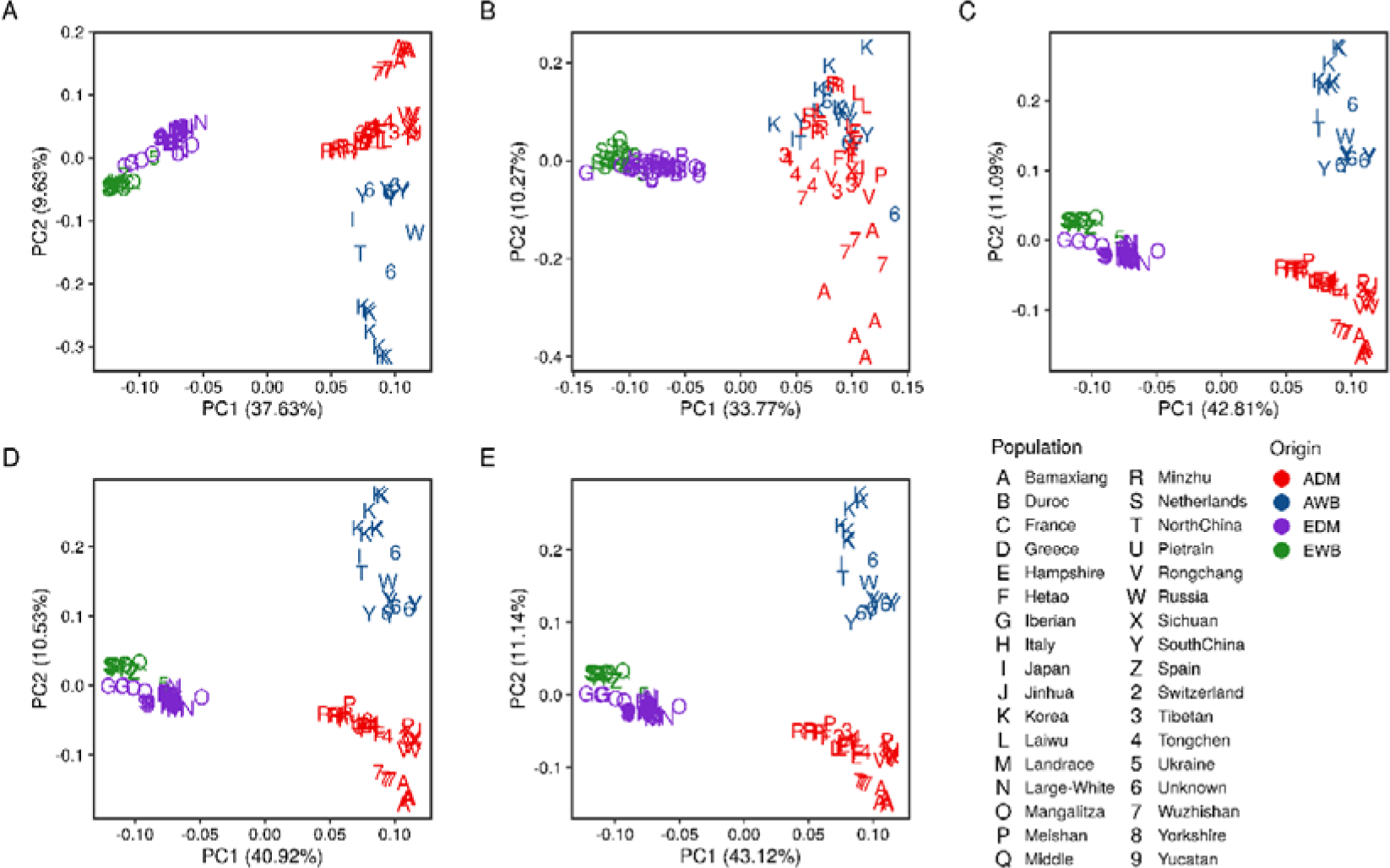
Principal component analysis plots based on SNPs mapping to: (**A**) the whole-genome (N = 19,720,314 SNPs), (**B**) miRNA genes (N = 285 SNPs), (**C**) Full 3’UTRs (N = 709,343 SNPs), (**D**) 3’UTR 7mer-m8 sites (N = 107,196 SNPs) and (**E**) 3’UTR 8mer sites (N = 33,511 SNPs), respectively.

With regard to miRNA variability, the 370 porcine miRNA genes annotated in the manually curated miRCarta v1.1 database [44] were selected, and SNPs within these genes were retrieved. A total of 285 SNPs residing in 139 miRNAs (37.56% of the total count) were identified (**Additional file 2: Table S2**), implying that most of miRNAs are monomorphic. The majority of these 139 miRNA loci (76.98%) contained 1-2 SNPs located inside their predicted genomic boundaries, while 18.70% contained between 3 up to 5 variants, and 4.32% of them displayed more than 7 SNPs (**Additional file 8: Fig. S2**). Only 43 miRNA SNPs (15.09%) were shared amongst EDM, ADM, EWB and AWB populations (**Additional file 2: Table S2**), showing alternative alleles in at least one of the analyzed individuals in each group. The number of SNPs segregating in each of the four defined groups were 129 (EDM), 201 (ADM), 76 (EWB) and 172 (AWB), respectively (**Additional file 2: Table S2**). With regard to precursor and mature regions, 41 and 2 SNPs were shared amongst the four populations under consideration, respectively (**Additional file 9: Fig. S3A** and **S3B**). None of the SNPs in the seed regions were shared by the four porcine populations (**Additional file 9: Fig. S3C**). Only three miRNA SNPs were found in the European data set but not in the Asian one. In strong contrast, 55 miRNA SNPs were detected in the Asian data set but not in the European one. Principal component analyses based on identified autosomal miRNA SNPs (N = 260, **Additional file 2: Table S2**) showed the existence of a poor differentiation between pigs and wild boars (**Fig. 2B**), while the genetic divergence between European and Asian individuals was still apparent.

When we analyzed the population structure based on whole-genome autosomal 3’UTR SNPs (N = 709,343 SNPs, **Fig. 2C**), 3’UTR 7mer-m8 site SNPs (N = 107,196 SNPs, **Fig. 2D**) and 3’UTR 8mer site SNPs (N = 33,511 SNPs, **Fig. 2E**), the genetic differentiation between Asian vs European populations was evident, in close concordance with results shown in **Fig. 2A** and **2B**. However, we also detected a more pronounced differentiation between domestic pigs and wild boars, and this observation was particularly applicable to Asian pigs and wild boars (**Fig. 2C-D**).

### The analysis of European and Asian populations shows reduced variability in porcine microRNAs

About 47.76%, 57.36%, 44.77% and 36.84% of miRNA SNPs showed alternative allele frequencies ≤ 0.1 in the ADM, EDM, AWB and EWB populations, respectively (**Fig. 3, Additional file 2: Table S2**). Variations located at mature miRNA and seed regions were enriched in rare or very rare variants when compared to the variability of miRNA precursor regions (**Additional file 2: Table S2**), with average alternative allele frequencies of ∼0.1 for mature and seed miRNA polymorphisms. In contrast, the average alternative allele frequency observed for SNPs in precursor areas was ∼0.15.

**Fig. 3:**
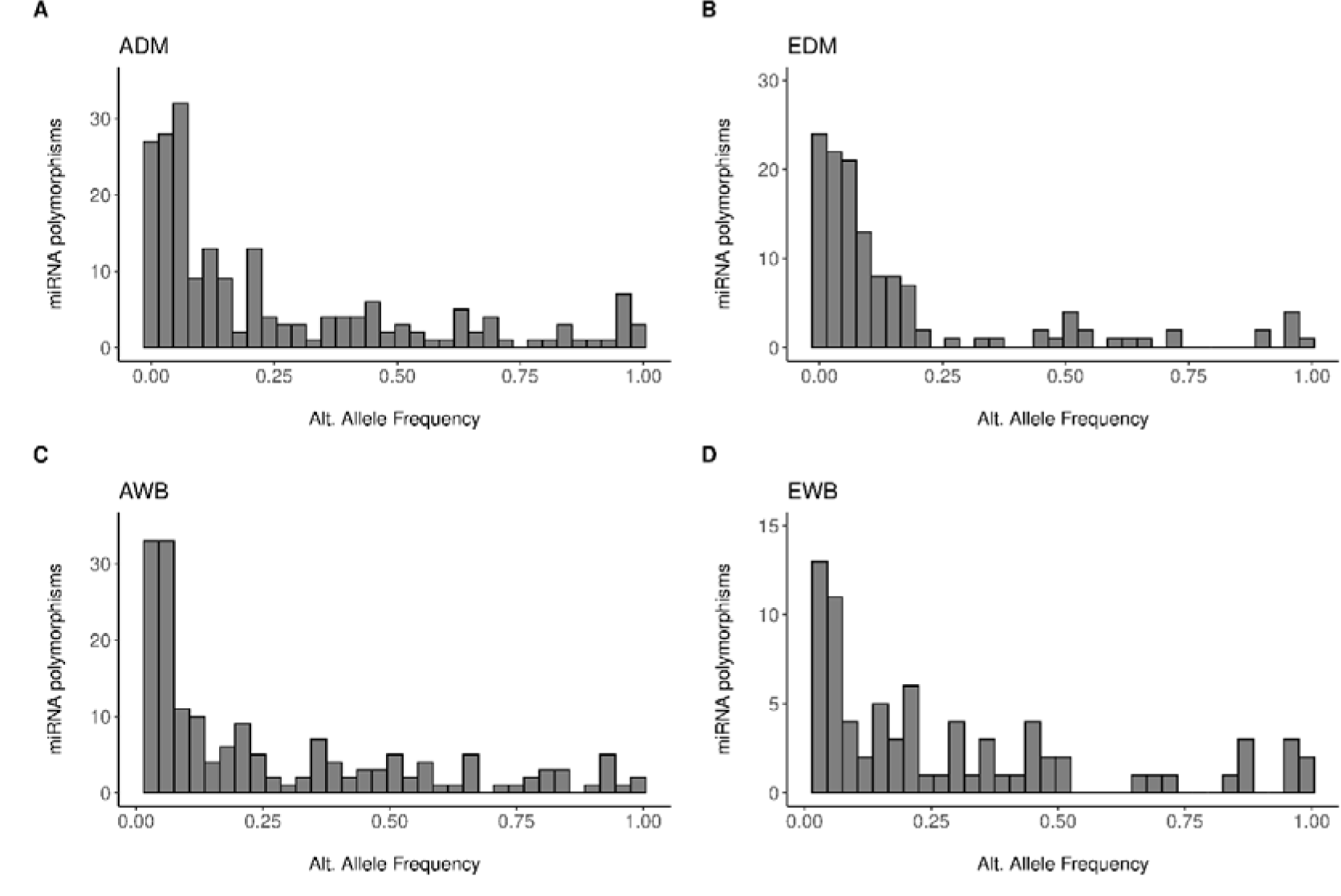
Alternative allele frequency distribution of polymorphisms located at miRNA loci in (**A**) Asian domestic pigs (ADM), (**B**) European domestic (EDM) pigs, (**C**) Asian wild boars (AWB) and (**D**) European wild boars (EWB).

Moreover, the observed SNP density adjusted to 100 bp for miRNA precursor, mature and seed regions consistently followed the order *precursor* > *mature* > *seed* when we considered the full set of 120 WGS. Indeed, ∼1 SNP per 100 bp was detected in precursor regions, whereas ∼0.86 and ∼0.42 SNPs per 100 bp were observed in the mature and seed regions, respectively (**Fig. 4A**). After computing a non-parametric comparison of SNP density between the mature and seed regions with a Mann Whitney U test [50], a significant difference (*P*-value = 0.016) was detected. These results implied a 1.16-fold reduction in SNP density between precursor and mature regions, while for seed regions, which are critical determinants of miRNA-mRNA interaction, the observed SNP density was ∼2.4 fold lower than in precursor regions. This differential distribution of the SNP density across miRNA regions (*precursor* > *mature* > *seed*) was also observed in each of the analyses performed in the ADM, EDM, AWB and EWB groups (**Fig. 4A**). With regard to variants located within mature miRNAs (N = 64), both inside (N = 12) and outside seed regions (N = 52), their distribution along the sequence of the mature miRNA (∼22 nts) showed a characteristic pattern (**Fig. 4B**): amongst all the detected SNPs, the 1^st^ position of the mature miRNA 5’ end, which binds to the MID domain of the Argonaute protein in the miRISC complex, exhibited a SNP density of ∼0.49 SNPs per 100 bp. Such scarcity in polymorphic sites was also observed when considering the next 2^nd^ to 8^th^ positions in the mature miRNA sequence (seed region), with an average of ∼0.42 SNPs per 100 bp across the whole seed and up to ∼0.73 SNPs per 100 bp in the 6th position of the mature miRNA. In contrast, the interval comprising positions 9^th^ to 12^th^ (with no functional implications in terms of mRNA targeting) showed an increased average SNP density of ∼0.98 SNPs per 100 bp. Positions 13^th^ to 18^th^ of the mature miRNA, which roughly correspond to a functional region providing additional anchor pairing to the seed region, showed a decreased SNP density, particularly at positions 16^th^ to 17^th^ (**Fig. 4B**). Additionally, an increased SNP density was found at positions 19^th^ to 22^nd^. Furthermore, the SNP density in miRNA genes (N = 370, **Fig. 4C**) was ∼2.6 fold lower (*P*-value < 0.01, see Methods) than in their flanking regions (± 1 kb). A list of the miRNA SNPs (N = 64) located at mature miRNAs and their genomic coordinates within the mature miRNA sequence is available at **Additional file 10: Table S7.**

**Fig. 4:**
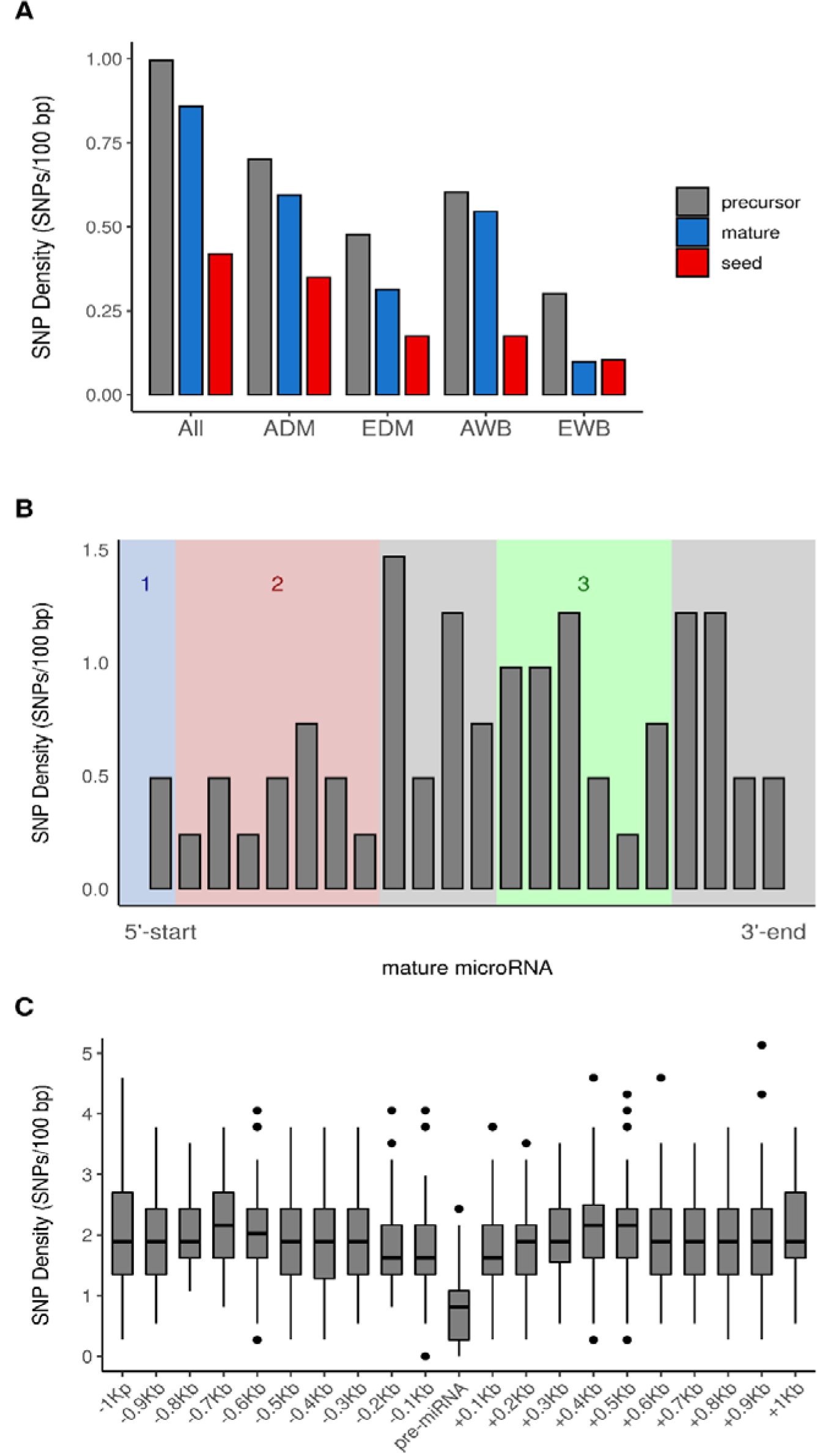
(**A**) SNP density per 100 bp for each analyzed miRNA region considering the full set of 120 porcine whole-genome sequences (WGS) as well as each WGS from the ADM, EDM, AWB and EWB groups. (**B**) SNP density across mature miRNA regions. 1: anchor (1^st^ 5’ end position), 2: seed (2^nd^ to 8^th^ position) and 3: supplemental pairing (13^th^ to 18^th^ position). (**C**) SNP density in miRNA loci and flanking regions considering a window of ± 1kb divided in 10 upstream and downstream 100 bp bins.

### Statistics of the whole-genome sequencing of five Duroc boars

As previously explained, we sequenced the genomes of five Duroc pigs which founded a population of 350 offspring with the goal of identifying SNPs in miRNA genes and investigating their association with mRNA expression and lipid phenotypes. Mean coverage values of the five pig genomes ranged from 37.67× to 46.6×, with more than 98.6% of the genome covered by at least 10 reads in all five samples, and 96.71% of the genome covered by at least 15 reads. More details about coverage and genome mapping parameters are shown in **Additional files 11** and **12: Table S8** and **Fig. S4**. After performing variant calling on mapped reads, a total of 13,839,422 SNPs passed the established quality filters, whereas 3,721,589 insertions and deletions (INDELs) were detected. Moreover, a total of 1,643,861 INDELs (44.17%) and 6,034,548 SNPs (43.60%) were located in annotated protein coding loci. A total of 55 SNPs and 5 INDELs were located in the 370 miRNA loci annotated in the Sscrofa.11.1 genome assembly according to the miRCarta database [44]. From these, 53 variants (96.36%) were shared with the set of 285 miRNA SNPs found in the 120 WGS reported before (**Additional file 13: Table S9**). After performing quality check of the variants mapping to miRNA genes, 49 SNPs and 5 INDELs passed all established filtering criteria.

Amongst the 49 identified miRNA polymorphisms, we selected 12 SNPs on the basis of their location at relevant annotated miRNA loci (**Table 1, Additional file 13: Table S9**) for genotyping and performing association analyses. Microarray expression data comprised a total of 23,998 probes, of which 12,131 (50.55%) and 12,787 (53.28%) corresponded to genes expressed in GM and liver tissues, respectively.

To provide a comprehensive view about the association between miRNA genotypes and liver and muscle mRNA levels, we have carried out an association analysis between miRNA genotypes and the whole set of expressed probes/genes in these two tissues (**Table 2**). It can be seen that there are hundreds of associations at the nominal level, while the number of significant associations is strongly reduced after multiple testing correction. This analysis does not take into account neither the existence of complementarity between the seed of the miRNA and the porcine 3’UTR of target mRNAs, nor functional validation data obtained in humans (please see the Methods section). A second analysis taking into account these two criteria revealed that from the set of 12 genotyped SNPs listed in **Table 1**, only 2 SNPs showed significant associations (*q*-value < 0.1) with the levels of mRNAs predicted to be miRNA targets (**Table 3, Additional file 14: Table S10**). Further details of target sites at the 3’UTR of mRNA transcript expression profiles significantly associated with miRNA SNP genotypes (**Table 3**) are available at **Additional file 15: Table S11**). All genes displaying significant associations (*q*-value < 0.1) in this second analysis ranked amongst the top 1% of significance (**Additional file 16: Table S12**) in the condition-free analysis (Analysis 1), except for transcripts associated with ssc-miR-326 in the liver, which ranked amongst the top 5% (**Table 1, Additional file 16: Table S12**). We observed that for several of the significant associations between miRNA and mRNA pairs reported in **Table 3** and **Additional file 14: Table S10**, the interaction was classified as 7-mer-m8 by our targeting approach with the *locate* tool of SeqKit toolkit [46] when, in reality, it was of type 8mer with one mismatch (data not shown).

**Table 2:** Number of significant associations between the 12 genotyped miRNA SNPs and the whole set of mRNA expressed in the *gluteus* (GM) skeletal muscle (N = 89) and liver (N = 87) tissue samples from Duroc pigs.

| SNP | Type | Significant Associations |  |  |  |
| --- | --- | --- | --- | --- | --- |
|  |  | GM (*P*-value < 0.05) | LIVER (*P*-value < 0.05) | GM (*q*-value < 0.1) | LIVER (*q*-value < 0.1) |
| rs81349391 (1:164026014) | ssc-miR-339 apical loop (A/G) | 456 | 356 | 0 | 0 |
| rs344472188 (2:13296736) | ssc-miR-130a apical loop (A/G) | 320 | 1252 | 0 | 1 |
| rs333787816 (2:65308181) | ssc-miR-23a precursor stem (T/C) | 2075 | 751 | 6 | 1 |
| rs340704946 (4:6948743) | ssc-miR-30d precursor stem (A/G) | 563 | 398 | 0 | 0 |
| rs320008166 (6:56427227) | ssc-miR-371 precursor stem (T/C) | 270 | 365 | 2 | 8 |
| rs323906663 (6:63491948) | ssc-miR-429 precursor stem (G/A) | 328 | 406 | 2 | 8 |
| rs319154814 (9:9581989) | ssc-miR-326 apical loop (G/A) | 372 | 2400 | 0 | 0 |
| rs321151601 (9:39280312) | ssc-miR-34c mature region (C/A) | 947 | 392 | 0 | 0 |
| rs341950320 (12:36947510) | ssc-miR-378-2 precursor stem (G/A) | 792 | 322 | 0 | 0 |
| rs334680106 (13:100083229) | ssc-miR-15b precursor stem (C/T) | 277 | 205 | 0 | 0 |
| rs335924546 (17:10758828) | ssc-miR-486 precursor stem (C/T) | 1232 | 171 | 0 | 0 |
| rs334590580 (18:18341582) | ssc-miR-335 precursor stem (T/C) | 665 | 219 | 1 | 0 |

**Table 3:**
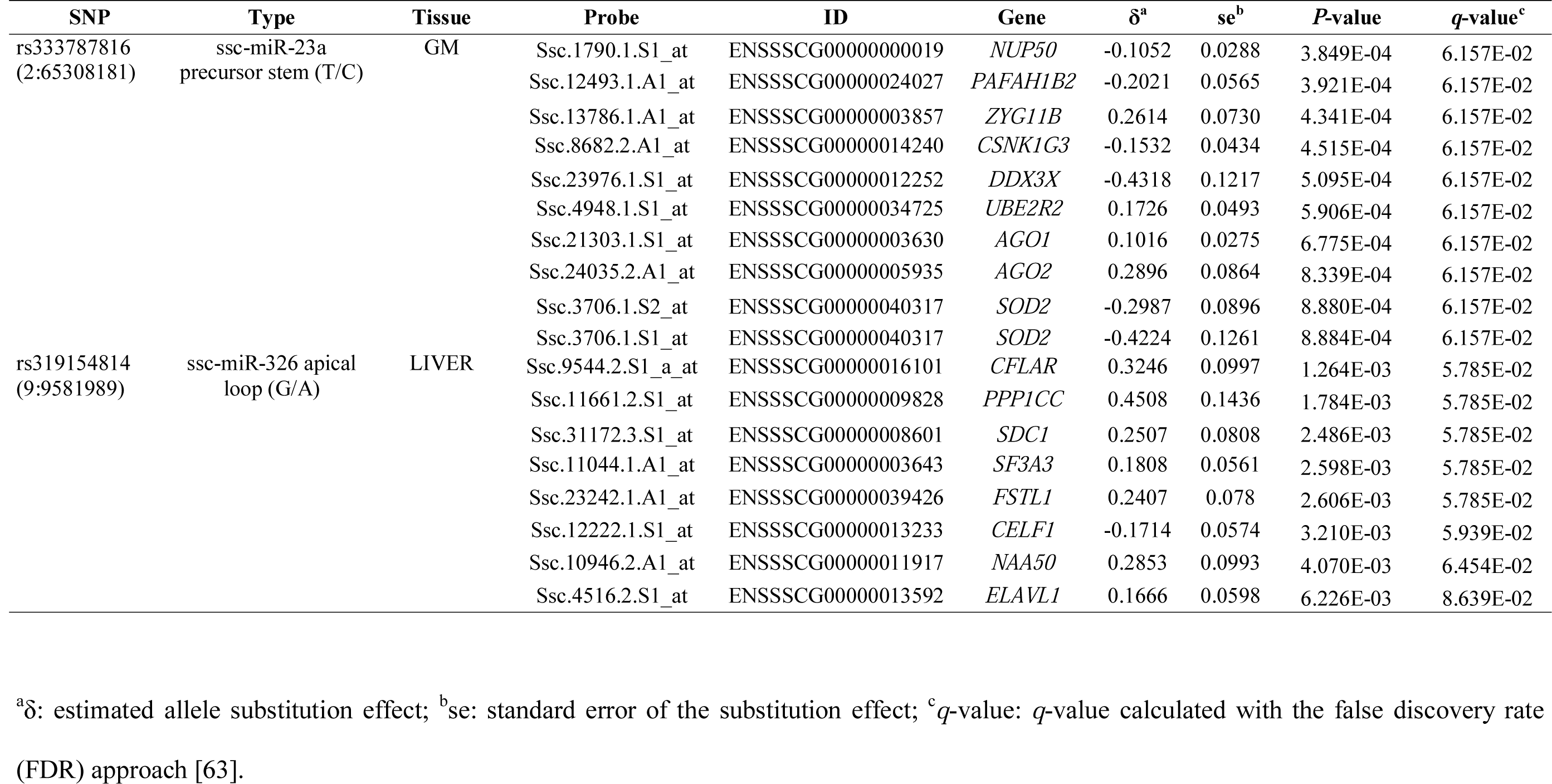
Top significant associations after correction for multiple testing (*q*-value < 0.1) between miRNA SNPs and the mRNA expression of otential targets in the *gluteus medius* (GM) skeletal muscle (N = 89) and liver (N = 87) tissue samples from Duroc pigs.

### Two SNPs in the apical loop of ssc-miR-326 and in the precursor region of ssc-miR-23a are associated with the mRNA expression of several of their putative targets

When we analyzed the association between the rs319154814 (G/A) polymorphism located in the apical loop of ssc-miR-326 and gene expression data (**Table 3**), significant results were obtained after multiple testing correction (*q*-value < 0.1). More specifically, we detected eight significant associations between this SNP and the hepatic mRNA expression of experimentally confirmed targets of this miRNA (**Table 3**). For instance, the hepatic mRNA levels of the cellular FLICE-like inhibitory protein (*CFLAR*), the protein phosphatase 1 catalytic subunit γ (*PPP1CC*), the syndecan 1 (*SDC1*), the splicing factor 3A subunit 3 (*SF3A3*) or the Follistatin-like 1 (*FSTL1*) were amongst the most significantly associated with ssc-miR-326 genotypes (**Table 3**). The expression levels of seven of these mRNAs associated with ssc-miR-326 rs319154814 (G/A) genotypes (**Table 3**) were reduced in pigs homozygous for the mutated allele (AA, N = 32), as depicted in **Fig. 5**.

**Fig. 5:**
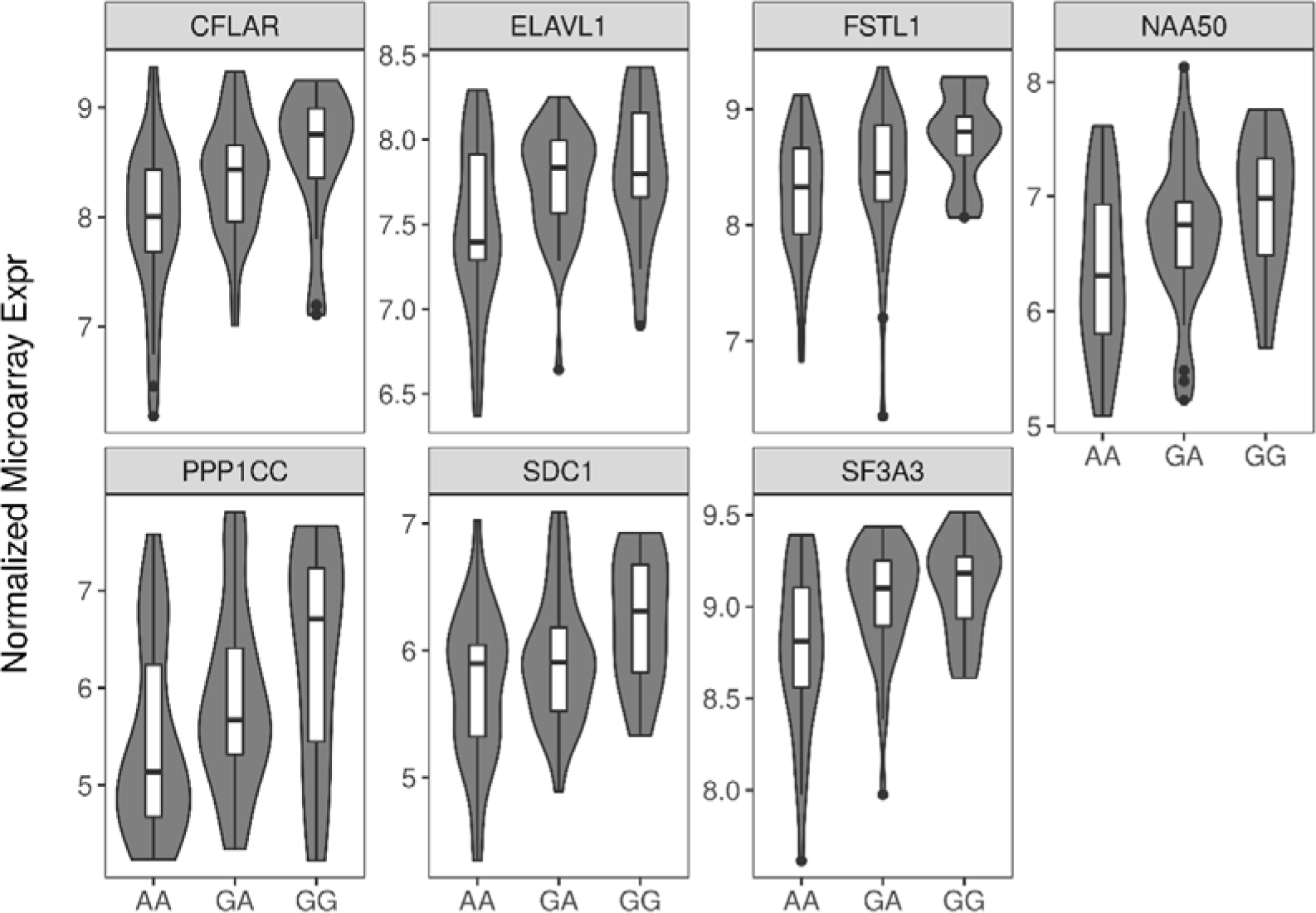
Hepatic mRNA expression levels of the *CFLAR*, *PPP1CC, SDC1, SF3A3, FSTL1*, *NAA50* and *ELAVL1* genes according to the genotype of the rs319154814 (G/A) apical loop SNP in the ssc-miR-326 gene. The number of individuals representing each genotype were: GG (N = 17), GA (N = 37) and AA (N = 32).

The mRNA levels of the *PPP1CC*, *CFLAR* and *SF3A3* mRNA transcripts were measured by RT-qPCR in 10 liver samples (5 belonging to each AA or GG rs319154814 genotypes). After comparing the mean expression for each of the three analyzed mRNA transcripts in both AA and GG genotype groups, only the *PPP1CC* transcripts showed a significantly reduced expression in AA pigs compared with their GG counterparts (*P*-value = 0.027, **Fig. 6A**). The other two transcripts (*CFLAR* and *SF3A3*) also showed a reduced expression in pigs with AA genotype (**Fig. 6A**), but results were not significant. We further investigated whether the ssc-miR-326 rs319154814 (G/A) genotypes were associated with the levels of this very same miRNA in the liver by using a specific Taqman Advanced miRNA probe assay. In this qPCR analysis, pigs with AA genotypes for the rs319154814 polymorphism showed an increased ∼1.9-fold expression of ssc-miR-326 transcripts compared with their GG counterparts, although such difference did not reach statistical significance (*P*-value = 0.178, **Fig. 6B**).

**Fig. 6:**
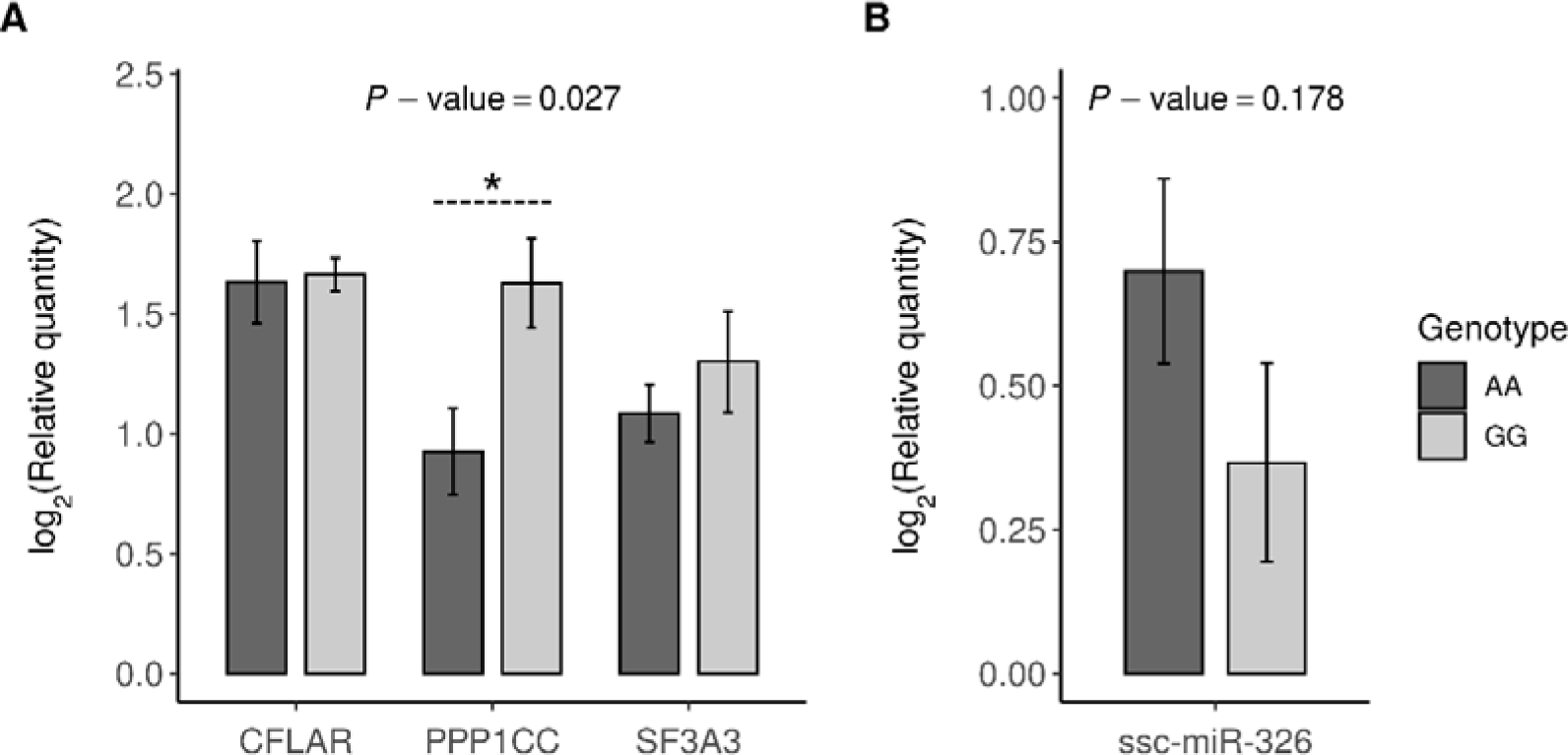
(**A**) Barplots depicting qPCR log_2_ transformed relative quantities (Rq) for *CFLAR, PPP1CC* and *SF3A3* mRNA transcripts measured in the liver tissue. The number of individuals representing each genotype were: GG (N = 5) and AA (N = 5). (**B**) Barplots depicting qPCR log_2_ transformed relative quantities (Rq) for ssc-miR-326 transcripts measured in the liver tissue. The number of individuals representing each genotype were: GG (N = 9) and AA (N = 9).

The rs333787816 (T/C) polymorphism, located in the precursor region, immediately downstream the mature ssc-miR-23a sequence, was also significantly associated with 36 experimentally confirmed targeted genes in the GM muscle tissue (*q*-value < 0.1, **Table 3, Additional file 14: Table S10**). From these, it is worth mentioning the Argonaute RISC component 1 (*AGO1*) and the Argonaute RISC catalytic component 2 (*AGO2*) mRNAs. Both *AGO1* and *AGO2* genes showed lower mRNA expression levels in homozygous CC pigs with respect to their TT and TC counterparts (**Table 3**).

### Pathway enrichment analyses of significantly associated mRNA targets provide insights of putative metabolic functions of miRNAs

After performing pathway enrichment analyses on the sets of putative targeted mRNA transcripts significantly associated with miRNA SNP genotypes (*P*-value < 0.05), several biological categories were identified and reported at **Additional file 17: Table S13**. To mention a few of the most relevant findings, the ssc-miR-23a genotype was associated with several regulatory pathways (post-transcriptional silencing by small RNAs, competing endogenous RNAs regulate PTEN translation, regulation of PTEN mRNA translation, regulation of RUNX1 Expression and Activity etc.) involving genes, such as *AGO1*, *AGO2* or *TNRC6C* genes, which play relevant roles in miRNA-mediated gene regulation [74]. Amongst the sets of mRNAs nominally associated with ssc-miR-30d genotype in GM, we identified the regulation of ornithine decarboxylase activity or cellular response to hypoxia, as well as mature mRNA transport after splicing (**Additional file 17: Table S13**).

In the liver tissue, the ssc-miR-326 genotype was significantly associated with the expression of several circadian clock regulatory transcripts e.g., histone deacetylase 3 (*HDAC3*), period 1 (*PER1*) and the already mentioned *PPP1CC* gene. Moreover, negative regulation of DDX58/IFIH1 signaling of type I interferon response and p53 signaling were pathways enriched in the sets of mRNAs nominally associated with ssc-miR-130a genotypes (**Additional file 17: Table S13**).

### Porcine lipid phenotypes are associated with the genotypes of miRNA genes

We also evaluated the association between miRNA SNPs and several lipid-related phenotypes recorded in the Lipgen population (**Additional file 3: Table S3**). Only the rs319154814 (G/A) variant in the ssc-miR-326 gene was significantly associated (*q*-value < 0.1) with lipid traits (**Table 4**, **Additional file 18: Table S14**). More specifically, we found significant associations with myristic acid (C14:0) content in both LD and GM muscles, as well as with the gadoleic acid (C20:1) content and the ratio between PUFA and MUFA in the LD muscle (**Table 4, Additional file 18: Table S14**). We also observed several associations between the rs319154814 (G/A) SNP and fatty acid composition traits, but they were significant only at the nominal level (*P*-value < 0.01, **Table 4**). Other apical loop SNPs like rs81349391 (A/G) at ssc-miR-339 and rs344472188 (T/C) at ssc-miR-130a were significantly associated at the nominal level (*P-*value < 0.01) with palmitic acid (C16:0) content and SFA and UFA proportion in the GM muscle, as well as with backfat thickness. As shown in **Table 4**, a SNP located in the precursor 3’ stem of ssc-miR-30d showed a nominally significant association with the content of arachidic acid (C20:0). Other significant associations at the nominal level were, for instance, those between the rs334590580 (T/C) SNP located at the precursor stem region of ssc-miR-335 and palmitic and arachidic acids content in GM tissue (**Table 4**).

**Table 4:**
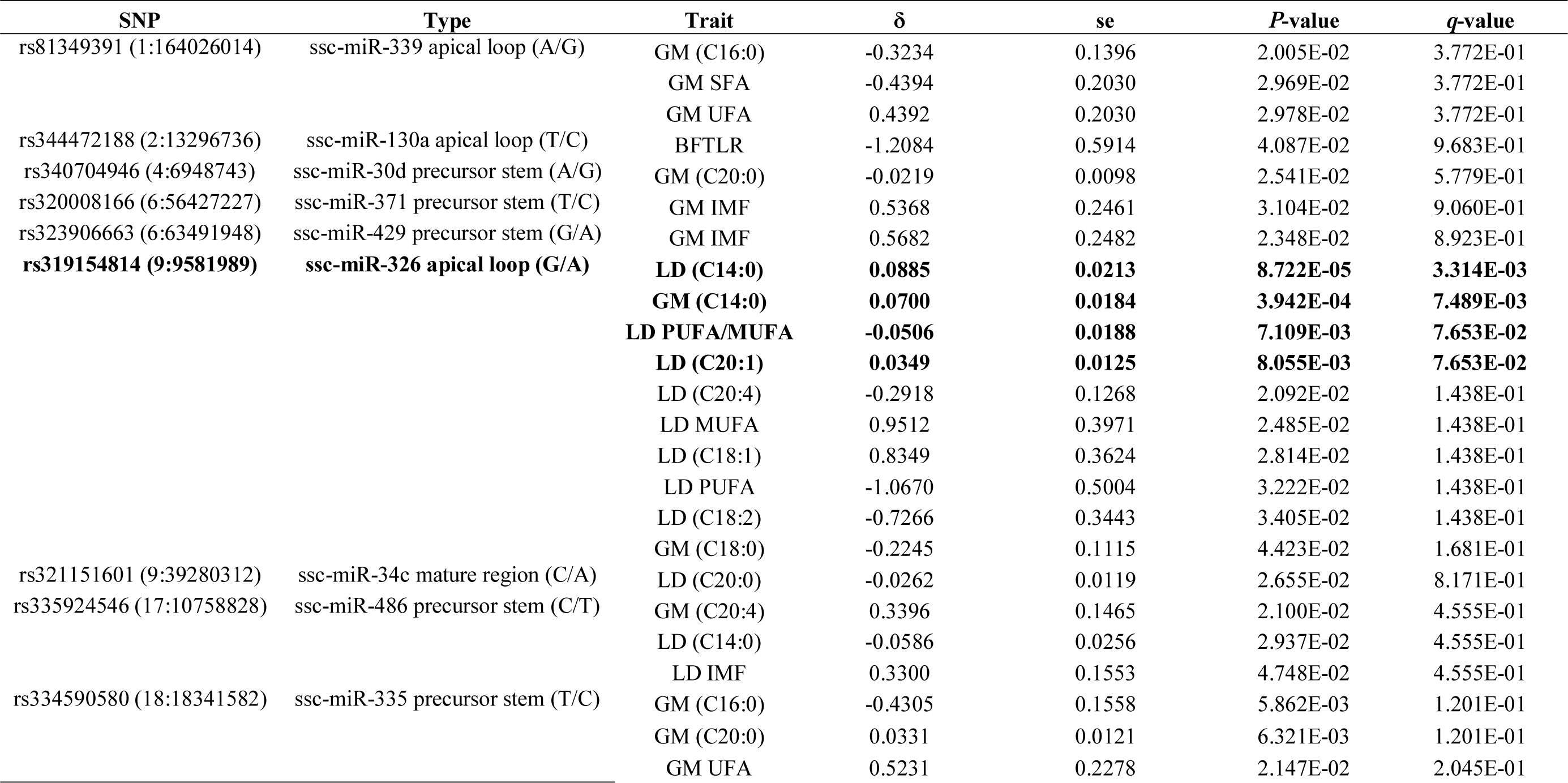
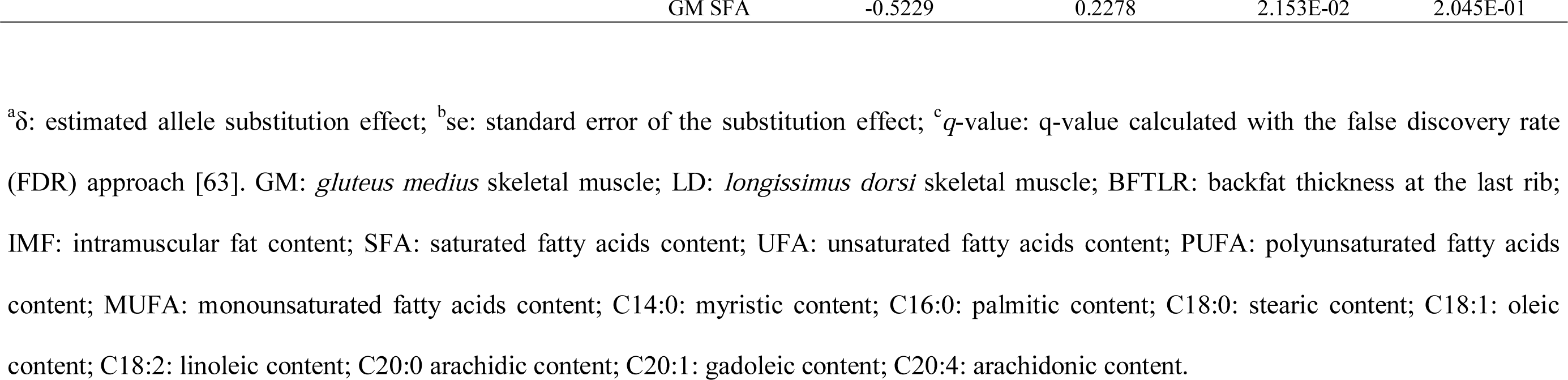
Significant associations at the nominal level (*P*-value < 0.01) and after multiple testing correction (*q*-value < 0.1; in bold) between 12 ped miRNA SNPs and lipid phenotypes recorded in a population of Duroc pigs (N = 345).

## Discussion

### Divergent patterns of variation for microRNA and 3’UTR polymorphisms in Asian and European pigs and wild boars

The PCA revealed the existence of a detectable genetic differentiation between Asian and European populations, with the latter showing reduced levels of diversity when compared to the former. Groenen et al. (2012) [75] investigated the variability of pig genomes and found that Asian pigs and wild boars are more diverse than their European counterparts and that both gene pools split during the mid-Pleistocene 1.6–0.8

Myr ago. Calabrian glacial intervals probably favored a restricted gene flow between these two gene pools [75]. The high variability of Asian populations could be explained by the fact that *Sus scrofa* emerged as a species in Southeast Asia (5.3-3.5 Mya) and then dispersed westwards until reaching Europe around 0.8 Mya [76]. This initial founder effect combined with the occurrence of strong bottlenecks reduced the genetic diversity of European wild boars [75].

While genetic differentiation between wild boar and pig populations was clearly discernible considering the whole-genome SNP data set (**Fig. 2A**), this was less evident in the PCA based on miRNA SNPs (**Fig. 2B**), probably because the low number (285 SNPs) of markers employed in this analysis limits the resolution with which population differentiation can be detected. We have also found that the degree of population differentiation between Asian domestic pigs and Asian wild boars increases when PCAs are built on the basis of SNPs located in the 3’UTR (709,343 SNPs), 3’UTR 7mer-m8 sites (107,196 SNPs) or 3’UTR 8mer (33,511 SNPs) sites (**Figs. 2C-E**). The potential effects of 3’UTR SNPs are the modulation of mRNA expression, secondary structure, stability, localization, translation, and binding to miRNAs and RNA-binding proteins [77], so in general, they are not expected to have drastic consequences on gene function [77]. Purifying selection is less intense in 3’UTRs than in protein-coding regions, implying that 3’UTRs evolve faster and accumulate a larger fraction of recent polymorphisms contributing to population differentiation [78].

### Low SNP density in microRNA genes and lack of uniform SNP distribution across sites

We have found that, in general, miRNA loci have a substantially lowered SNP density in their seeds when compared with mature and precursor regions (**Fig. 4A**). When we analyzed the SNP density in miRNA loci and their flanking regions (± 1 kb), a significantly reduced amount of SNPs per 100 bp was found in miRNAs compared with upstream and downstream flanking sequences (**Fig. 4C**). These results were in agreement with data reported by Omariba et al. (2020) [79] and Saunders et al. (2007) [5].

The low variability of miRNA genes, a feature that was particularly evident in the seed region, is probably due to the intense effects of purifying selection. Indeed, the importance of the miRNA seeds is revealed by the high conservation of their sequence across species [16,23,80], as this sequence ultimately determines the success of miRNA-mRNA interactions [2]. In our study, a total of 221, 52 and 12 SNPs were found in precursor, mature and seed regions within miRNA loci, respectively (**Additional files 2** and **9: Table S2, Fig. S3**). Gong et al. (2012) [81] described the existence of 40% polymorphic miRNAs in the human genome but only 16% of them displayed more than one SNP. In a more recent study, He et al. (2018) [82] reported 1,879 SNPs in 1226 (43.6%) human miRNA seed regions, and 97.5% of these polymorphisms had frequencies below 5%. These results agree well with the overall frequency distribution of miRNA SNPs in the European and Asian populations analyzed in the current work (**Fig. 3**). He et al. (2018) also demonstrated that 1,587, 749, 340, 102, 31, and 4 miRNAs harbored zero, one, two, three, four, and five SNPs, respectively, in their seed regions, reflecting that mutations in this critical functional region are not well tolerated [82]. This distribution is similar to the one that we have observed in domestic pigs and wild boars, with 81, 31, 11, 9 and 5 miRNAs harboring one, two, three, four and five SNPs (**Additional file 8: Fig. S2**). Only 4 and 2 miRNAs showed a total of seven and ten polymorphisms within their sequences (**Additional file 2: Table S2**).

We have also detected a non-uniform SNP density along the sequence of mature miRNAs (**Fig. 4B**). Gong et al. (2012) [81] showed that SNPs tend to concentrate in the middle region of the mature miRNA gene rather than in its 5’ and 3’ends. Moreover, the same authors described an increased SNP density at positions 9 and 15 of the mature miRNA, a result that agrees well with ours (**Fig. 4B**) However, we have also identified an elevated number of polymorphic sites at positions 11, 19 and 20, a finding that does not match human data presented by Gong et al. (2012) [81].

Several of the sites showing a reduced variability in porcine miRNAs exert critical functions (**Fig. 4B**, **Additional file 10: Table S7**). For instance, the 1^st^ nucleotide of mature miRNAs plays an important role in the loading process of the mature miRNA on the Argonaute protein to form the miRISC complex [83]. Nucleotides 2^nd^ to 8^th^ in the mature miRNA correspond to the seed, where we found a consistently reduced SNP density (**Fig. 4A** and **4B**) compared with other miRNA regions. This result was expected because this region has a crucial role in modulating the interaction between the mature miRNA and its 3’UTR binding sites. Polymorphisms in the seed region have the potential to disrupt the proper miRNA-mRNA pairing and thus alter biologically relevant regulatory pathways, which tend to be evolutionarily conserved [80]. Hence, these variants located in the miRNA seeds (**Additional file 10: Table S7**) might favor the emergence of novel miRNA-mRNA interactions as well as the abolishment of conserved ones, thus modifying gene regulatory networks.

In contrast with the first eight 5’ positions of the miRNA, we found a higher SNP density in positions 9^th^ to 12^th^, which do not contribute substantially to miRNA target recognition [2] (**Fig. 4B**). Positions 13^th^ to 16^th^ facilitate 3’-compensatory pairing between the mature miRNA and targeted 3’UTRs [84], although only at marginal levels [85]. Nevertheless, in our porcine data set only positions 16^th^-17^th^ showed a reduced SNP density (**Fig. 4B**).

### Polymorphisms in microRNA genes show associations with the mRNA expression of several of their predicted targets

We have investigated the association of 12 SNPs mapping to miRNA genes and hepatic and muscle mRNA expression in Duroc pigs. Several hundreds of significant associations at the nominal level (*P*-value < 0.05) were detected in both GM muscle and liver tissues when considering the whole set of microarray expression profiles of mRNA transcripts (**Table 2**). One pitfall of this analysis is that it does not take into account whether there is molecular evidence supporting the existence of direct miRNA-mRNA interactions displaying significant associations. On the other hand, indirect interactions between miRNAs and mRNAs might also exist i.e., a SNP regulating the expression of a miRNA might repress the translation of a given mRNA through a direct interaction (an event that would not necessarily imply any transcriptional decay, hence being undetectable in our experimental system) and, in turn, this might indirectly affect the expression of other multiple mRNAs regulated by the translationally repressed mRNA. Despite this consideration, we decided to restrict our association analyses to miRNA-mRNA pairs with molecular evidence of potential interactions because we believe that with this stringent approach, we are able to minimize the chances of detecting spurious false positive results. The genomic position of miRNA-binding sequences in the 3’UTR of targeted mRNAs showing significant associations (*q*-value < 0.1, **Table 3**) with miRNA genotypes is available in **Additional file 15: Table S11**).

After correction for multiple testing (*q*-value < 0.1, **Table 3**), only SNPs mapping to the ssc-miR-23a and ssc-miR-326 genes displayed significant associations with the expression levels of some of their putative mRNA targets. More in detail, the rs333787816 (T/C) polymorphism in the ssc-miR-23a gene showed significant associations with several putative target mRNAs, amongst which are *AGO1* and *AGO2* transcripts, two essential components of the miRNA-mediated silencing machinery [86, 87]. The rs319154814 (G/A) polymorphism in the apical loop region of ssc-miR-326 also showed a significant association, after multiple testing correction, with the hepatic mRNA expression of several of its predicted and experimentally confirmed targets (**Table 3**). In contrast, no association with GM muscle mRNA expression was observed. Tissue-specific differences in the expression of the miRNA or its mRNA targets might explain such outcome. Indeed, the analysis of the distribution of miRNA expression across human tissues has shown that only a minority of miRNAs are expressed ubiquitously [88]. The hepatic mRNA targets showing the most significant association with rs319154814 (G/A) ssc-miR-326 genotype were *CFLAR*, *PPP1CC, SDC1, SF3A3* and *FSTL1*. The protein encoded by the *PPP1CC* gene belongs to the protein phosphatase PP1 subfamily, which is a ubiquitous serine/threonine phosphatase involved in regulating multiple cellular processes through dephosphorylation signaling. Amongst them, it is worth mentioning insulin signaling [89], post-translational localization of circadian clock components [90] and lipids [91, 92] or glycogen metabolism regulation [93]. Moreover, the cellular FLICE-like inhibitory protein (*CFLAR*) gene encodes the cFLIP protein and is involved in the inhibition of Fas-mediated apoptosis [94], while the Follistatin-like 1 (*FSTL1*) gene plays a role in the immune inflammatory signaling and fibrosis in the liver [95].

Pigs homozygous for the derived A-allele of the rs319154814 SNP showed a consistent downregulation of the hepatic mRNA expression of the *CFLAR*, *PPP1CC, SDC1, SF3A3, FSTL1, NAA50* and *ELAVL1* genes (**Fig. 5**), a result that was confirmed by qPCR for *PPP1CC* mRNA transcripts (*P*-value = 0.027, **Fig. 6A**). Although in the qPCR experiment the *CFLAR* and *SF3A3* genes also showed decreased mRNA levels in pigs with homozygous AA genotypes for the rs319154814 SNP, such associations did not reach statistical significance (**Fig. 6A**). The significant downregulation of at least one top mRNA putative target (i.e., *PPP1CC*) might suggest that the rs319154814 (G/A) variant could increase the repressive activity of ssc-miR-326. Indeed, when we assessed, with a dedicated qPCR assay, the expression level of ssc-miR-326 transcripts in the liver tissue, we found that ssc-miR-326 log_2_ relative expression is increased by ∼1.9 fold in pigs homozygous for the A-allele, although such result was not statistically significant (*P*-value = 0.178, **Fig. 6B**).

As previously discussed, the rs319154814 (G/A) polymorphism is located in the apical loop region of ssc-miR-326. Although the apical region of the miRNA does not have a function as critical as the seed, polymorphisms located in this specific region can have relevant effects on the structural conformation of the pre-miRNA hairpin. Indeed, Fernandez et al. (2017) [19] described a mutation in the apical loop of hsa-miR-30c (G/A) that induces a steric disruption of the pri-miRNA folding structure of the hairpin, hence creating a bulge around the flanking downstream CNNC motif that facilitates the SRSF3 factor accessibility to the RNA sequence [96]. In other words, SNPs in the apical region can modify the efficiency with which the Drosha processing machinery, is recruited. The rs319154814 (G/A) polymorphism detected in our study might have structural consequences similar to those described for the hsa-miR-30c apical loop variant [19].

One clear limitation of our experimental design is that we did not test the existence of miRNA-mRNA interactions by using functional assays. Such experiments would be necessary to demonstrate our hypothesis that the rs319154814 (G/A) polymorphism in the apical loop of ssc-miR-326 has a role in promoting the processing and maturation efficiency of the hairpin precursor, thus modifying the expression profiles of ssc-miR-326 and its putative mRNA targets.

### A polymorphism in the apical loop of microRNA 326 is associated with fatty acid composition traits

In our study, the only miRNA polymorphism showing significant associations (*q*-value < 0.1) with lipid-related phenotypes was the rs319154814 (G/A) SNP in the apical loop of ssc-miR-326. This SNP was associated with myristic fatty acid content in the GM and LD muscles of Duroc pigs, as well as with PUFA/MUFA ratio or gadoleic fatty acid content in LD. To the best of our knowledge, no direct effect of ssc-miR-326 on the metabolism of myristic fatty acid has been described so far, but there are reports suggesting that several of the targets of this miRNA might be involved in carbohydrate and lipid metabolism pathways [97, 98]. Moreover, the *PPP1CC* transcript, one of the predicted targets of miR-326, encodes a subunit of protein phosphatase-1, which activates acetyl-CoA carboxylase α (ACACA) and 6-phosphofructo2-kinase/fructose-2,6-bisphosphatase (PFKFB), the main regulators of fatty acid synthesis and glycolysis, respectively [92]. The protein phosphatase-1 also activates lipogenic transcription factors such as sterol regulatory element-binding protein 1 (SREBF1), carbohydrate-responsive element-binding protein (MLXIPL) and, moreover, it dephosphorylates the DNA-dependent protein kinase encoded by the *PRKDC* gene, which is another main determinant of hepatic lipogenesis [92]. In summary, a potential effect of the rs319154814 (G/A) SNP in the synthesis or degradation of myristic acid can be envisaged, but this hypothesis still needs to be confirmed at the functional level. Interestingly, Cardoso et al. (2021) [99] explored the relationship between miRNA SNPs located in the seed and mature regions of bovine miRNA loci and fatty acid composition traits. By combining lipid-related phenotype data with mRNA and protein abundances in a multi-omics approach, they described three miRNA SNPs significantly associated with several unsaturated and polyunsaturated fatty acid traits, as well as with the expression levels of some of their predicted targets [99].

### Conclusions

MicroRNA genes show divergent patterns of variation between Asian and European pigs and wild boars and, in general, they display low levels of polymorphism. As expected, this reduced miRNA variability was particularly prevalent in the seed region, a finding that is likely explained by the strong effects of purifying selection aiming to preserve the conservation of this critical site. We have detected one SNP in the apical loop of ssc-miR-326 and another one in the precursor region of ssc-miR-23a that are associated with the mRNA expression of several of their putative targets. If confirmed with functional assays, these results would reinforce the need of exploring the role of miRNA variation within and outside the seed in the fine-tuning of mRNA expression in pigs.

## Declarations

### Ethics approval

Animal care and management procedures were carried out in accordance with national guidelines for Good Experimental Practices and they were approved by the Ethical Committee of the Institut de Recerca i Tecnologia Agroalimentàries (IRTA).

### Consent of publication

Not applicable.

### Availability of data materials

Microarray expression data used in the current study were deposited in the Gene Expression Omnibus (GEO) public repository and are accessible through GEO Series Accession Number GSE115484. Phenotypic and genotypic data sets generated and analyzed during the current study have been deposited in the Figshare public repository available at https://figshare.com/projects/SNPs_miRNA/78690. Whole-genome sequencing dataset from the 5 Duroc boars is available at the Sequence Read Archive (SRA) database (BioProject: PRJNA626370). Variant Calling Format (VCF) files from the 120 European and Asian pigs belonging to domestic breeds and wild boars, as well as from the 5 sequenced Duroc boars are available at the following link: https://figshare.com/projects/VCF_PIGs/93140.

### Authors’ contributions

MA conceived this study. MA and RQ designed the experimental protocols. RQ coordinated phenotyping recording and contributed to generate microarray expression data. EMS did DNA extractions and selected SNPs to be genotyped. EMS performed all bioinformatic and statistical analyses of the data. MGL and AC conducted qPCR analyses. DG contributed to population structure analyses. RT performed whole-genome sequencing of the five Duroc boars. EMS and MA drafted the manuscript. All authors read and approved the content of the final manuscript.

## Additional Files legends

**Additional file 1: Table S1:** List of whole-genome sequences from European domestic pigs (EDM, N = 40), Asian domestic pigs (ADM, N = 40), European wild boars (EWB, N = 20) and Asian wild boars (AWB, N = 20).

**Additional file 2: Table S2:** Single nucleotide polymorphisms located in microRNA genes and their frequencies in European (E) and Asian (A) domestic pigs (DM) and wild boars (WB).

**Additional file 3: Table S3**: Means and standard deviations (SD) of fatness and intramuscular fat content and composition traits recorded in the *gluteus medius* (GM) and *longissimus dorsi* (LD) muscles of 345 Duroc pigs.

**Additional file 4: Fig. S1:** Sequence homology between 12 porcine miRNAs harboring SNPs that have been genotyped in the Lipgen population (N = 345) and their corresponding human miRNA orthologous sequences.

**Additional file 5: Table S4:** Set of primers used for the qPCR quantification of three mRNAs putatively targeted by ssc-miR-326.

**Additional file 6: Table S5:** Distribution of SNPs mapping to specific genomic annotated regions (i.e., protein coding genes, exons, introns, 3’UTRs, miRNAs, lncRNAs, snRNAs, snoRNAs and pseudogenes) and segregating in the set of whole-genome sequences from 120 Asian and European domestic pigs and wild boars.

**Additional file 7: Table S6:** List of porcine genes showing 3 or more multi-allelic SNPs in the set of 120 whole-genome sequences from Asian and European domestic pigs and wild boars.

**Additional file 8: Fig. S2:** Number of SNPs present in each of the annotated porcine miRNA loci that display at least 1 SNP.

**Additional file 9: Fig. S3:** Venn Diagrams illustrating the sharing of SNPs located at the (**A**) *precursor*, (**B**) *mature* and (**C**) *seed* regions of miRNA loci amongst Asian domestic pigs (ADM), Asian wild boars (AWB), European domestic pigs (EDM) and European wild boars (EWB).

**Additional file 10: Table S7:** Description of SNPs mapping to porcine mature miRNA loci (N = 409) and segregating in the set of 120 whole-genome sequences from Asian and European domestic pigs and wild boars.

**Additional file 11: Table S8:** Whole-genome sequencing statistics for the five Duroc boars that sired the Lipgen population (N = 345).

**Additional file 12: Fig. S4:** Boxplot distribution depicting whole-genome sequencing statistics for the five Duroc boars that sired the Lipgen population (N = 345).

**Additional file 13: Table S9:** List of SNPs mapping to 370 porcine miRNA loci and segregating in the 5 Duroc boars that sired the Lipgen population (N = 345).

**Additional file 14: Table S10:** Association analyses between SNPs in miRNA genes and the mRNA levels of their predicted targets in the *gluteus medius* (GM) skeletal muscle and liver tissues of Lipgen pigs.

**Additional file 15: Table S11:** Genomic coordinates and sequence of the seed (miRNA) and its predicted 3’UTR binding site (mRNA) for putative miRNA-mRNA pairs showing significant associations after multiple testing correction (*q*-value < 0.1).

**Additional file 16: Table S12:** Association analyses between miRNA SNPs and the mRNA expression profiles of the whole set of expressed probes/genes in the porcine *gluteus medius* (GM) skeletal muscle and liver tissues from Lipgen pigs.

**Additional file 17: Table S13:** Pathway enrichment analyses of the lists of probes/genes significantly associated at the nominal level (*P*-value < 0.05) with miRNA SNP genotypes.

**Additional file 18: Table S14:** Association analyses between miRNA SNPs and fatness and intramuscular fat content and composition traits recorded in the *gluteus medius* (GM) and *longissimus dorsi* (LD) skeletal muscles of 345 Duroc pigs.

**Additional file 4: Fig. S1:**
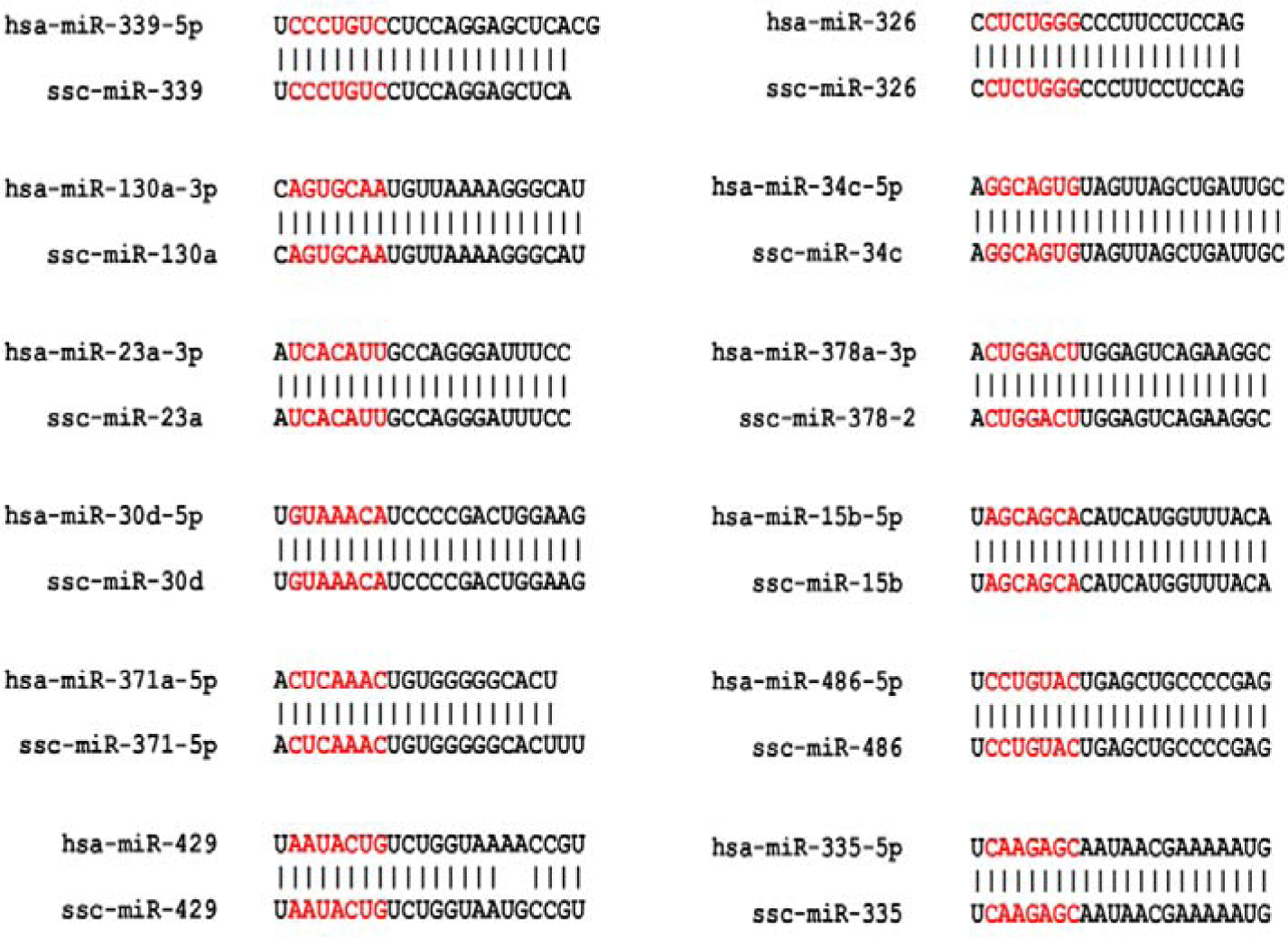
Sequence homology between 12 porcine miRNAs harboring SNPs that have been genotyped in the Lipgen population (N = 345) and their corresponding human miRNA orthologous sequences.

**Additional file 8: Fig. S2:**
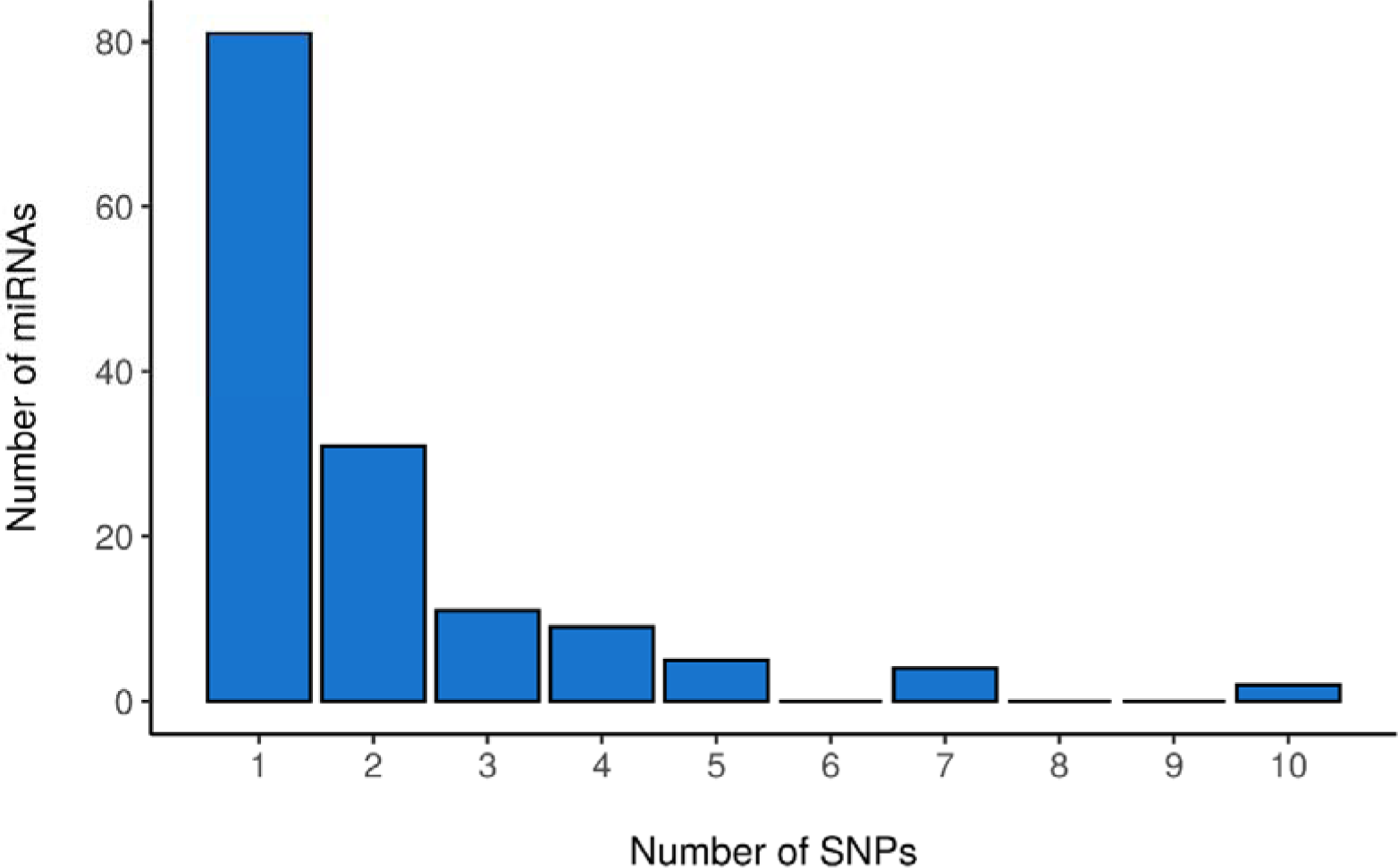
Number of SNPs present in each of the annotated porcine miRNA loci that display at least **1** SNP.

**Additional file 9: Fig. S3:**
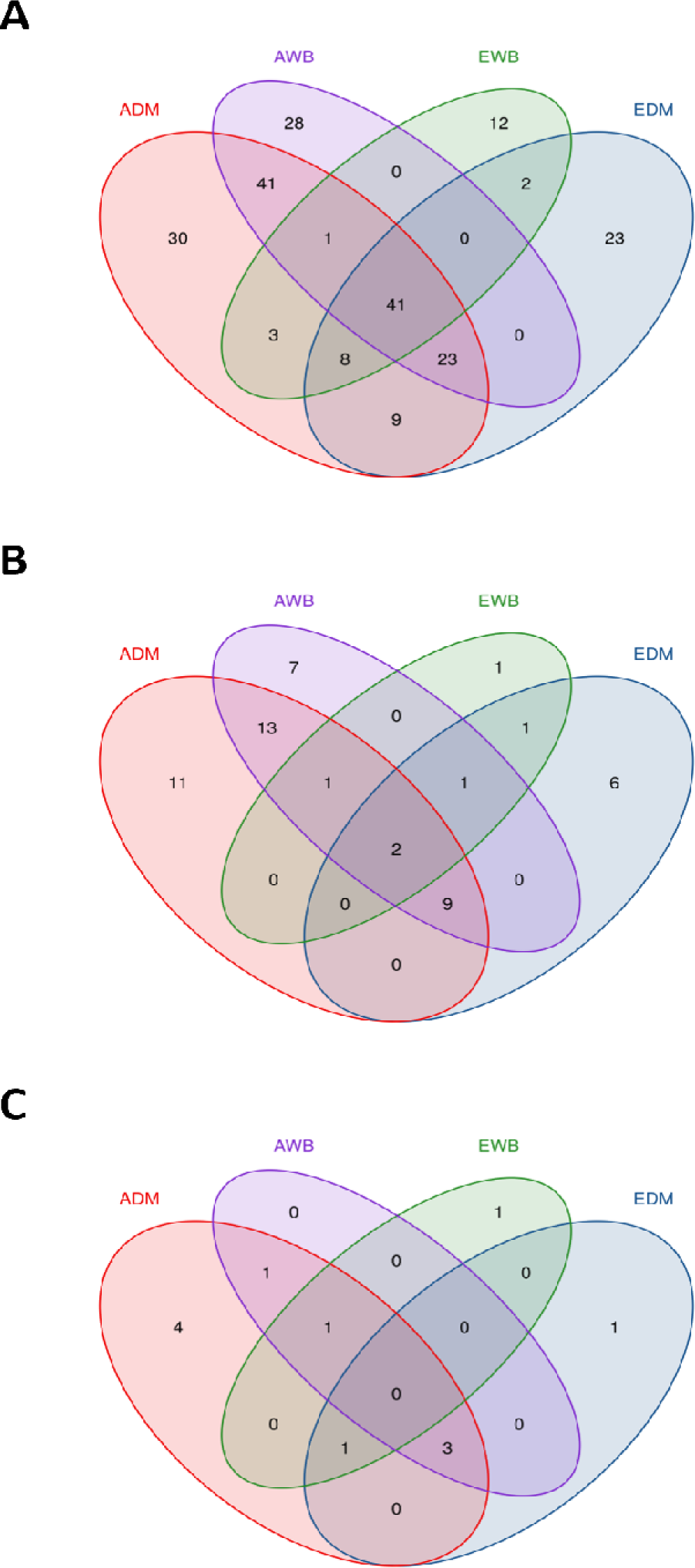
Venn Diagrams illustrating the sharing of SNPs located at the (A) *precursor,* **(B)** *mature* and (C) *seed* regions of miRNA loci amongst Asian domestic pigs (ADM), Asian wild boars (AWB), European domestic pigs (EDM) and European wild boars (EWB).

**Additional file 12: Fig. S4:**
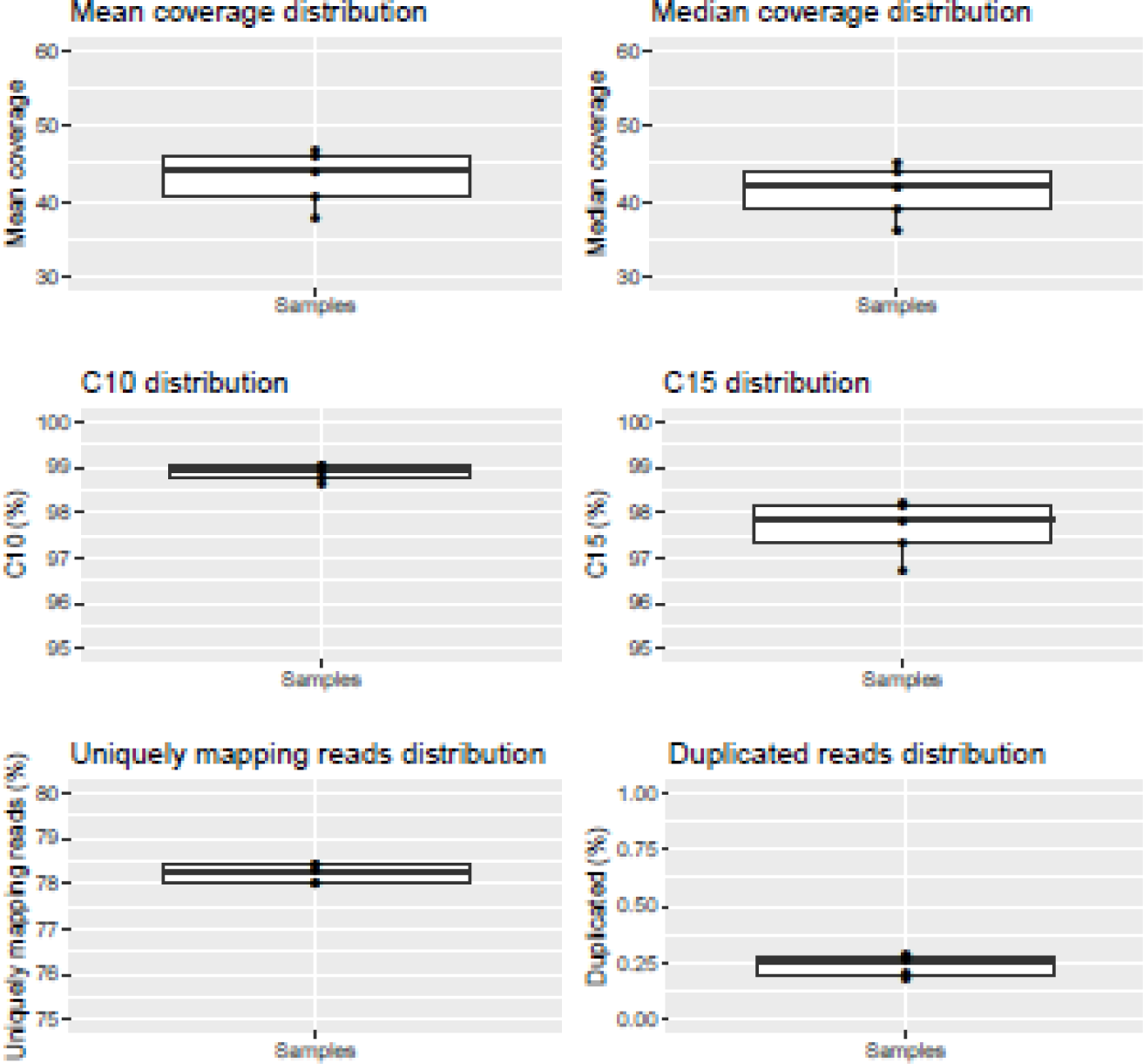
Boxplot distribution depicting whole-genome sequencing statistics for the five Duroc boars that sired the Lipgen population (N = 345).

